# Tradeoffs in Modeling Context Dependency in Complex Trait Genetics

**DOI:** 10.1101/2023.06.21.545998

**Authors:** Eric Weine, Samuel Pattillo Smith, Rebecca Kathryn Knowlton, Arbel Harpak

## Abstract

Genetic effects on complex traits may depend on context, such as age, sex, environmental exposures or social settings. However, it is often unclear if the extent of context dependency, or Gene-by-Environment interaction (GxE), merits more involved models than the additive model typically used to analyze data from genome-wide association studies (GWAS). Here, we suggest considering the utility of GxE models in GWAS as a tradeoff between bias and variance parameters. In particular, We derive a decision rule for choosing between competing models for the estimation of allelic effects. The rule weighs the increased estimation noise when context is considered against the potential bias when context dependency is ignored. In the empirical example of GxSex in human physiology, the increased noise of context-specific estimation often outweighs the bias reduction, rendering GxE models less useful when variants are considered independently. However, we argue that for complex traits, the joint consideration of context dependency across many variants mitigates both noise and bias. As a result, polygenic GxE models can improve both estimation and trait prediction. Finally, we exemplify (using GxDiet effects on longevity in fruit flies) how analyses based on independently ascertained “top hits” alone can be misleading, and that considering polygenic patterns of GxE can improve interpretation.

## Introduction

In organisms and study systems where the environment can be tractably manipulated, gene-by-environment interactions (GxE) are the rule, not the exception [1–5]. Yet, in complex (polygenic) human traits, there are but a few cases in which models that incorporate GxE explain data—such as Genome-Wide Association Study (GWAS) data—better than parsimonious models that assume additive contributions of genetic and environmental factors [6–8]. This is true for both physical environments but also for other definitions of “E,” broadly construed to be any context that modifies genetic effects, such as age, sex, or social setting [9–16].

GWAS commonly estimate marginal additive effects of an allele on a trait. The estimand here can be thought of as the average effect of the allele over a distribution of multidimensional contexts [17]. With this view, some differences in allelic effects across contexts are likely omnipresent, but may very well be small, such that the cost of including additional parameters (for context-specific effects) outweighs the benefit of measuring heterogeneous effects.

Here, we consider this problem and its connection to the currently underwhelming utility of GxE models in GWAS. First, we rigorously describe the statistical trade-off involved in estimating context-specificity at the level of a single variant. Then, we highlight ways in which this trade-off might change as we consider GxE in complex traits, involving numerous genetic variants simultaneously.

We begin by framing the problem of estimating context-specificity at an individual variant as a bias–variance trade-off. For example, consider the estimation of an allelic effect on lung cancer risk that depends on smoking status. When the allelic effect is estimated from a sample without considering smoking status, the estimate would be biased with respect to the true effect in smokers. We can estimate the effect separately in smokers and non-smokers to eliminate the bias, but the consideration of the additional parameters—smoking status-specific effects—has an associated cost of increasing the estimation variance, compared to an estimator that ignores smoking status. This bias–variance trade-off is closely related to the “signal-to-noise” ratio, where the signal of interest is the true difference in context-specific allelic effects. To demonstrate this tradeoff in real data, we consider sex-specific effects on physiological traits in humans. We show that for the majority of traits, it is typically unhelpful to model sex dependency for individual sites since the increase in noise vastly outweighs the signal.

We then consider the extension to GxE in complex traits. Complex trait variation is primarily due to numerous genetic variants of small effects distributed throughout the genome [18–21]. Simultaneously considering GxE across multiple variants may decrease estimation noise if the extent and mode of context-specificity is similar across numerous variants. This would tilt the scale in favor of context-dependent estimation. In addition, we show how conventional approaches for detecting and characterizing GxE, which focus on the most significant associations, may lead to erroneous conclusions. Finally, we discuss implications for complex trait prediction (with polygenic scores). We suggest a future focus on prediction methods that empirically learn the extent and nature of context dependency by simultaneously considering GxE across many variants.

## Results and Discussion

### Modeling context-dependent effect estimation as a bias–variance trade-off

#### The problem setup

We consider a sample of *n* + *m* individuals characterized as being in one of two contexts, *A* or *B. n* of the individuals are in context *A* with the remaining *m* individuals in context *B*. We measure a continuous trait for each individual, denoted by

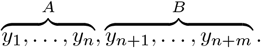

We begin by considering the estimation of the effect of a single variant on the continuous trait. We assume a generative model of the form

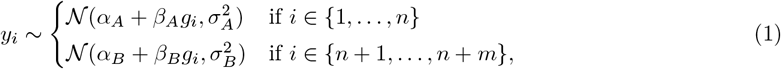

where *β*_*A*_ and *β*_*B*_ are fixed, context-specific effects of a reference allele at a biallelic, autosomal variant *i, g*_*i*_ ∈ {0, 1, 2} is the observed reference allele count. *α*_*A*_ and *α*_*B*_ are the context-specific intercepts, corresponding to the mean trait for individuals with zero reference alleles in context *A* and *B*, respectively. 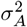 and 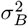 are context-specific observation variances. We would like to estimate the allelic effects *β*_*A*_ and *β*_*B*_.

#### Estimation approaches

We compare two approaches to this estimation problem. The first approach, which we refer to as GxE estimation, is to stratify the sample by context and separately perform an ordinary least squares (OLS) regression in each sample. This approach yields two estimates, 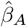 and 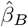,the OLS estimates of *β*_*A*_ and *β*_*B*_ of the generative model in **Eq. 1**, respectively. This estimation model is equivalent to a linear model with a term for the interaction between context and reference allele count, in the sense that context-specific allelic effect estimators have the same maximum likelihood estimators in the two models (see **Supplementary Materials**).

The second approach, which we refer to as additive estimation, is to perform an OLS regression on the entire sample and use the allelic effect estimate to estimate both *β*_*A*_ and *β*_*B*_. We denote this estimator as 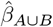,to emphasize that the regression is run on all individuals from context *A* and context *B*. This estimation model posits that for *i* = 1, …, *n* + *m*,

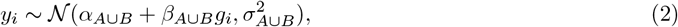

where *α*_*A*∪*B*_ is the mean trait value for an individual with zero reference alleles, *β*_*A*∪*B*_ is the additive allelic effect and 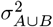 is the observation variance which is independent of context. Notably, this model differs from the generative model assumed above: *β*_*A*∪*B*_ may not equal *β*_*A*_ and *β*_*B*_; in addition, this model ignores heteroskedasticity across contexts.

#### Error analysis

We focus on the mean squared error (MSE) of the additive and GxE estimators for the allelic effect in context *A*. The estimator minimizing the MSE may differ between contexts A and B, but the analysis for context *B* is analogous. When selecting between these two estimation approaches, a bias–variance decomposition of the MSE is useful.

Based on OLS theory ([22, Theorem 11.3.3]), under the model specified above we have

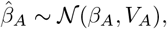

where 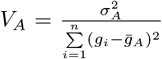 and 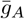 is the mean genotype of individuals in context *A*. The unbiasedness of the GxE estimator implies

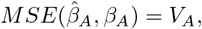

where 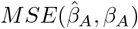 is the mean squared error of estimating *β*_*A*_ with 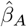.The case of the additive estimator, 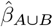,is a bit more involved. As we show in the **Methods** section, we can write

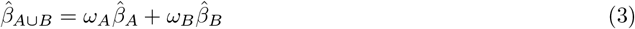

for non-negative weights *ω*_*A*_ and *ω*_*B*_ (that need not sum to 1). Further, we show in **Eq. 7** of the **Methods** section that *ω*_*A*_ ∝ *nH*_*A*_ and *ω*_*B*_ ∝ *mH*_*B*_, where *H*_*A*_ and *H*_*B*_ are the sample heterozygozities in contexts *A* and *B*, respectively. Using **Eq. 3**, we may write

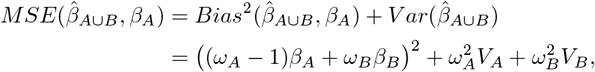

where *V*_*B*_ is defined analogously to *V*_*A*_. Thus, with MSE as our metric for comparison, we prefer the GxE estimator in context *A* when

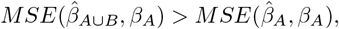

or, if and only if

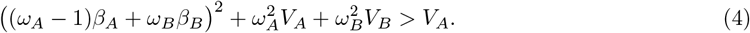

We refer to **Eq. 4** as the “decision rule,” since it guides us on the more accurate estimator; to minimize the MSE, we will use the context-specific estimator if and only if the inequality is satisfied.

To gain some intuition about the important parameters here, we first consider the case of equal allele frequencies (and hence equal heterozygozities) in both contexts and equal estimation variance in both contexts. In this case, the GxE estimator is advantaged by larger context-specificity (larger |*β*_*A*_ − *β*_*B*_|) and disadvantaged by larger estimation noise (larger *V*_*A*_ = *V*_*B*_) (**Fig. 1**). In fact, the decision boundary (i.e. the point at which the two models have equal MSE) can be written as a linear combination of |*β*_*A*_ − *β*_*B*_| and 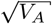 (**Fig. 1C**). In this special case, we show in the **Methods** section that **Eq. 4** is an equality when

**Figure 1.**
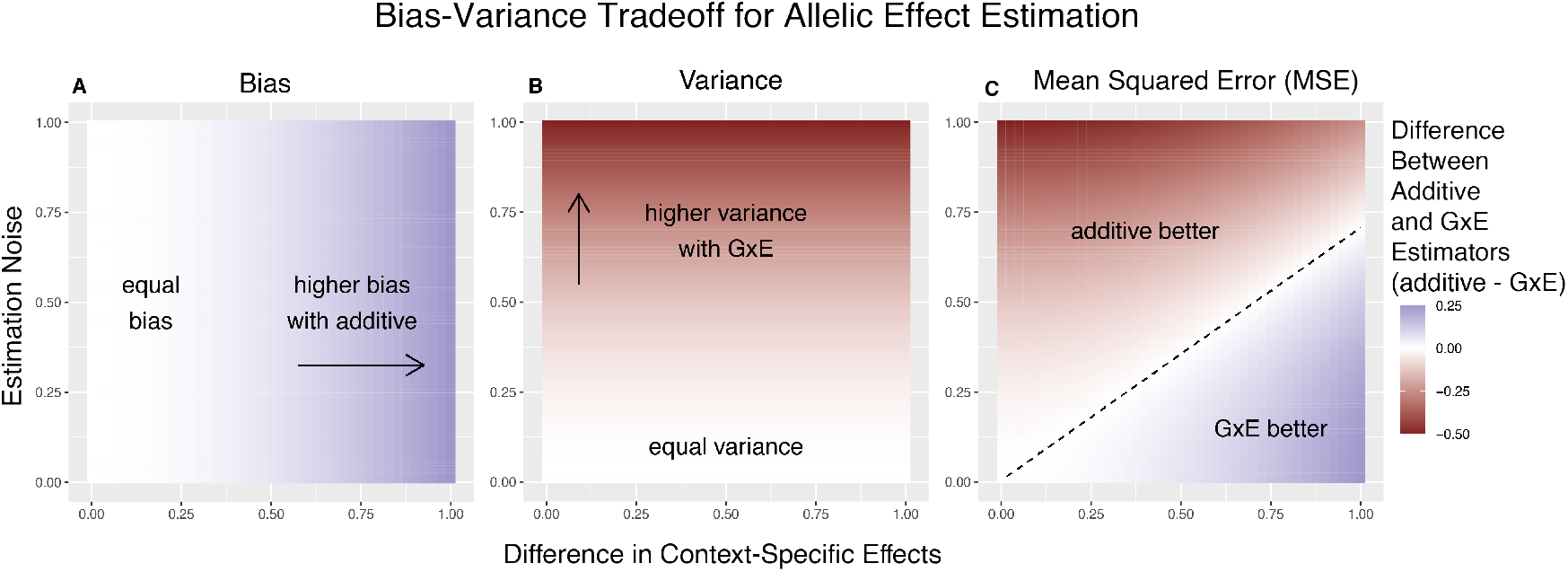
Bias-Variance tradeoff for single-site estimation with equal estimation noise and equal heterozygozity across contexts. The x-axis shows the difference in context-specific effects, while the y-axis shows the standard deviation of the context-specific estimators—both in raw measurement units. The color on the plot indicates the difference between the additive and GxE estimators in bias (A), variance (B) or MSE (C). **(A)** Only the additive estimator is potentially biased. The bias is proportional to the difference in contextspecific effects and independent of the estimation noise. **(B)** The difference in variance is proportional to context-specific estimation noise and independent of the difference of context-specific effects. **(C)** The decision boundary is linear in both the estimation noise and the difference between context-specific effects.

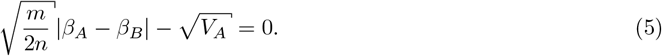

More generally, in the case where *H*_*A*_ = *H*_*B*_ but *V*_*A*_ ≠ *V*_*B*_, we show in the **Methods** section that we can write **Eq. 4** as

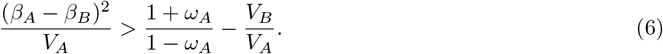

This dimensionless re-parameterization of the decision rule makes explicit its dependence on three factors. 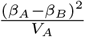 can be viewed as the “signal-to-noise” ratio: it captures the degree of context-specificity (the signal) relative to the estimation noise in the focal context, 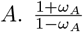 is the relative contribution to heterozygosity, which equals the relative contribution to variance in the independent variable of the OLS regression of **Eq. 2**. 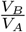 is the ratio of context-specific estimation noises. In the **Supplementary Materials**, we extend the decision rule for the case of a continuous context variable.

For a given trait and context, we can consider the behavior of the decision rule across variants with variable allele frequencies and allelic effects. The ratio of estimation noises, 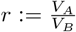,will not be constant. However, in some cases, considering a fixed *r* across variants is a good approximation. In GWAS of complex traits, each variant often explains a small fraction of trait variance. As a result, the estimation noise is effectively a matter of trait variance and heterozygosity alone. If per-site heterozygosity is similar in strata *A* and *B*, as it is, for example, for autosomal variants in biological males and females, *r* is approximately fixed across variants [9].

**Fig. 2** illustrates the linearity of the decision boundary under the assumption that *r* is fixed across variants. It also shows that the slope of the decision boundary changes as a function of *r*. Intuitively, we are less likely to prefer GxE estimation for the noisier context. In fact, for sufficiently small values of *r* (e.g. 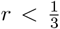 for 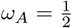), 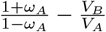 will be negative. This corresponds to the situation where *V*_*A*_ ≪ *V*_*B*_, in which case the additive estimator is never preferable to the GxE estimator in estimating *β*_*A*_, as the signal-to-noise ratio is always non-negative. Typically, this will also imply that the additive estimator is greatly preferable for estimating *β*_*B*_, as 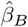 will be extremely noisy.

**Figure 2.**
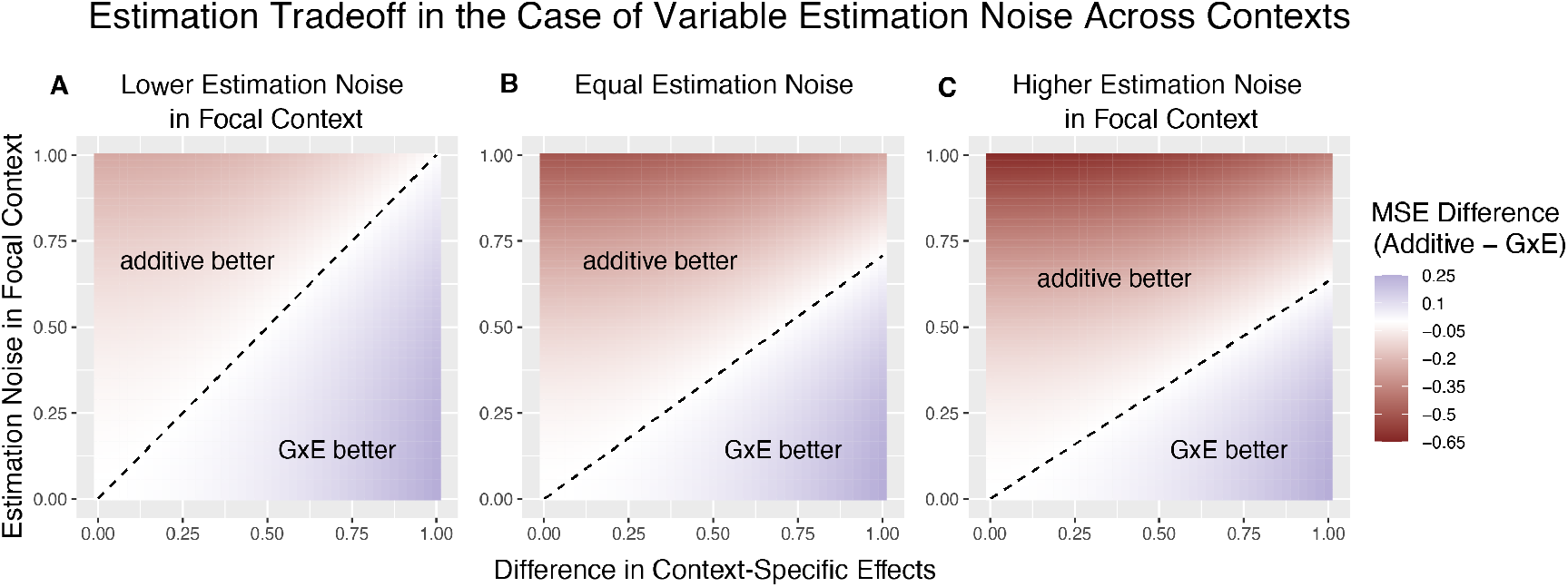
The decision boundary with different ratios of context-specific estimation noises. In all panels, the heterozygozity of the variant is assumed to be equal across contexts. The x and y axes are the same as in Fig. 1. Estimation noise in the focal context, A, is half that of the other context, B. **(B)** Estimation noise is equal in both contexts. **(C)** Estimation noise in focal context is double that of the other context.

It is natural to ask where the decision rule of **Eq. 4** falls with respect to empirical GWAS data. We considered the example of biological sex as the context (GxSex), and examined sex-stratified GWAS data across 27 continuous physiological traits in the UK Biobank [9, 23]. For each of nine million variants, we estimated the difference in sex-specific effects and the variance of each marginal effect estimator in males. Then, using an estimate of the ratio of sex-specific trait variances as a proxy for the ratio of estimation variances of males and females, we approximated the linear decision boundary between the additive and GxE estimators (**Fig. 3A,B**; **Text S1**). To demonstrate the accuracy of our decision rule, we employed a data-splitting technique where we estimate the MSE difference between estimators in a training set and evaluate the accuracy in a holdout set (**Fig. S1**).

**Figure 3.**
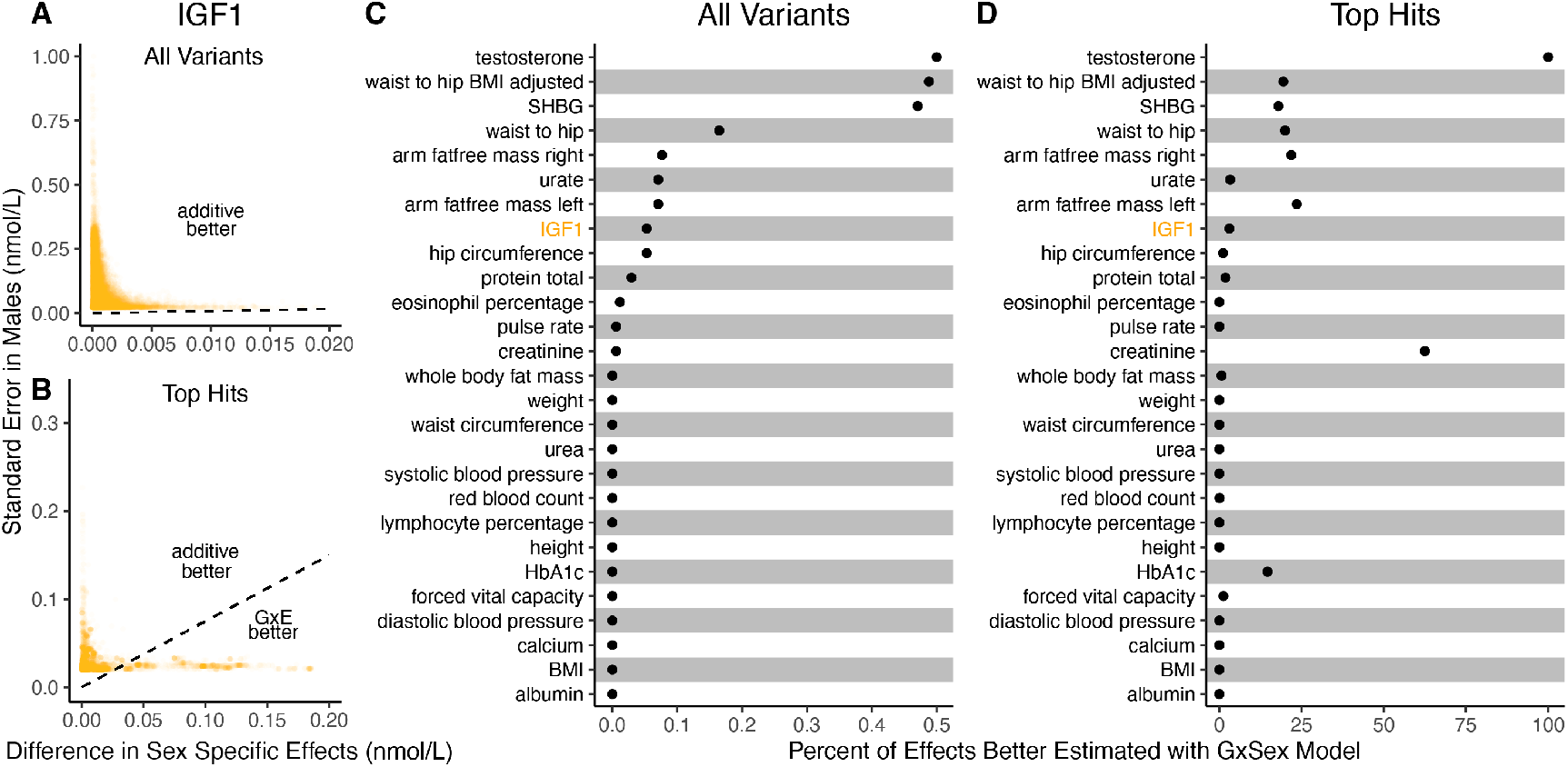
Applying the decision rule to sex-dependent effects on human physiological traits. **(A-B)** The x-axis shows the estimated absolute difference between the effect of variants in males and females. The y-axis shows the measured standard error for each variant in males, the focal context here. The dashed line shows the decision boundary for effect estimation in males. The difference in MSE between estimation methods increases linearly with distance from the dashed line, as in Fig. 2. If a variant falls above (below) the line, the additive (GxE) estimator has a lower MSE. (A) shows a random sample of 15K single nucleotide variants whereas (B) shows only variants with a marginal p-value less than 5 × 10^−8^ in males. **(C-D)** The percent of effects in males which would be better estimated by the GxE estimator, across continuous physiological traits. To estimate these percentages, one single nucleotide variant is sampled from each of 1,700 approximately independent autosomal linkage blocks, and this procedure is repeated 10 times. Shown are average percentages across the 10 iterations.

For almost all traits examined, very few allelic effects in males are expected to be more accurately estimated using the male-specific estimator (usually between 0% and 0.1%). Notable exceptions to this rule are testosterone, sex hormone binding globulin (SHBG), and waist-to-hip ratio adjusted for body mass index (BMI), for which roughly .5% of allelic effects are expected to be better estimated with the GxE model (**Fig. 3B**). However, when considering only SNPs that are genome-wide significant in males (marginal *p*-value < 5 × 10^−8^ in males), many traits show a much larger proportion of effects that would be better estimated by the GxE model. At an extreme, for testosterone, all genome-wide significant SNPs are expected to be better estimated by the GxE model. In addition, a large fraction of genome-wide significant effects are better estimated with the GxE model for creatinine (62%), arm fat-free mass (24%), waist-to-hip ratio (19%) and SHBG (18%) as well (**Fig. 3,D**).

The decision rule we derived could potentially guide more accurate allelic effect estimation approaches. However, the consideration of GxE pattern sharing across many variants (polygenic GxE) can alter both bias and variance and therefore the tradeoff. In our discussion of complex traits that follows, we therefore expand on the rule through qualitative consequences of polygenic GxE, and no longer stick to the analytical single variant rule.

### Context dependency in complex traits

At the single variant level, and specifically when variants are considered independently from one another, we have discussed how the accurate estimation of allelic effects can be boiled down to a bias–variance tradeoff. For complex traits, genetic variance is often dominated by the contribution of numerous variants of small effects that are best understood when analyzed jointly [8, 20, 24–28]. It stands to reason that to evaluate context-dependence in complex traits, we would also want to jointly consider polygenic patterns, rather than just the patterns at the loci most strongly associated with a trait [3, 13, 29–32].

Motivated by this rationale, we recently inferred polygenic GxSex patterns in human physiology [9]. One pattern that emerged as a common mode of GxSex across complex physiological traits is “amplification”: a systematic difference in the magnitude of genetic effects between the sexes. Moving beyond sex and considering any context, amplification can happen if, for example, many variants regulate a shared pathway that is moderated by a factor—and that factor varies in its distribution among contexts. Amplification is but one possible mode of polygenic GxE, but can serve as a guiding example for ways in which GxE may be pervasive but difficult to characterize with existing approaches [9, 16, 33, 34]. In what follows, we will therefore use the example of pervasive amplification (across causal effects) to illustrate the interpretive advantage of considering context dependency across variants jointly, rather than independently.

### A focus on “top hits” may lead to mis-characterization of polygenic GxE

A common approach to the analysis of context dependency involves two steps. First, categorization of context dependency (or lack thereof) is performed for each variant independently. Second, variants falling under each category are counted and annotated across the genome. Some recent examples of this approach towards the characterization of GxE in complex traits include studies of GxSex effects on flight performance in *Drosophila* [35], GxSex effects on various traits in humans [36, 37] and GxDietxAge effects on body weight in mice [38].

Characterizing polygenic trends by summarizing many independent hypothesis tests may miss GxE signals that are subtle and statistically undetectable at each individual variant, yet pervasive and substantial cumulatively across the genome. To characterize polygenic GxE based on just the “top hits” may lead to ascertainment biases, with respect to both the pervasiveness and the mode of GxE across the genome. Much like the heritability of complex traits is thought to be due to the contribution of many small (typically sub-significant) effects [24, 26], when GxE is pervasive we may expect that the sum of many small differences in context-specific effects accounts for the majority of GxE variation.

For concreteness, we consider in more depth one recent study characterizing GxDiet effects on longevity in *Drosophila melanogaster* [39]. In this study, Pallares et al. tracked caged fly populations given one of two diets: a “control” diet and a “high-sugar” diet. Across 271K single nucleotide variants, the authors tested for association between alleles and their survival to a sampling point (thought of as a proxy for “lifespan” or “longevity”) under each diet independently. Then, they classified variants according to whether or not their associations with survivorship were significant under each diet as follows:

1. significant under neither diet → classify as *no effect*.
2. significant when fed the high-sugar diet, but not when fed the control diet → classify as *high-sugar specific effect*.
3. significant when fed the control diet, but not when fed the high-sugar diet → classify as *control specific effect*.
4. significant under both diets → classify as *shared effect*.

This authors’ choice of four categories a variant may fall into may be motivated by the wish to test for the presence of “cryptic genetic variation”—genetic variation that is maintained in a context where it is functionally neutral but carries large effects in a new or stressful context [3, 5, 33, 40]. Indeed, of the variants Pallares et al. classified as having an effect (one hundredth of variants tested), approximately 31% were high-sugar specific, while the remaining 69% of the variants were shared. Fewer than 1% were labelled as having control specific effects. They concluded that high-sugar specific effects on longevity are pervasive, compatible with the hypothesis of widespread cryptic genetic variation for longevity.

This characterization of GxE, based on “top hits”, places an emphasis on the context(s) in which trait associations are statistically significant, rather than on estimating how the context-specific effects covary. In addition, this particular classification system also does not cover all possible ways in which context-specific effects may differ. In the **Supplementary Materials**, we discuss these interpretation difficulties further.

We next show that a generative model that differs qualitatively from the cryptic genetic variation model yields results that are highly similar to those observed by Pallares et al. We simulated data under pervasive amplification. Specifically, we sampled from a mixture of 40% of variants having no effect under either diet and 60% of variants having an effect under both diets—but exactly 1.4× larger under a high-sugar diet. We then simulated the noisy estimation of these effects, and employed the classification approach of Pallares et al. to the simulated data (**Methods**).

The patterns of allelic effects in the control compared to high-sugar contexts were qualitatively similar in the experimental data and our pervasive amplification simulation. This is true both genome-wide (**Fig. 4A** compared to **Fig. 4B**) and for the set of variants classified as significant with their classification approach (**Fig. 4C** compared to **Fig. 4D**). The similarity of ascertained variants further highlights caveats of interpretation based on the classification of “top hits”: despite the fact that we did not simulate any variants that only have an effect under the high-sugar diet, approximately 36% of significant variants were classified as specific to the high-sugar diet (green points in **Fig. 4D**), comparable to the 31% of variants classified as high-sugar specific in the experimental data (**Fig. 4C**). These variants simply have sub-significant associations in the control group and significant associations in the high-sugar group. In addition, every variant in the shared category (blue points in **Fig. 4D**) in fact has a larger effect in the high-sugar diet than in the control diet, which cannot be captured by the classification system itself but represents the only mode of GxE in our simulation.

**Figure 4.**
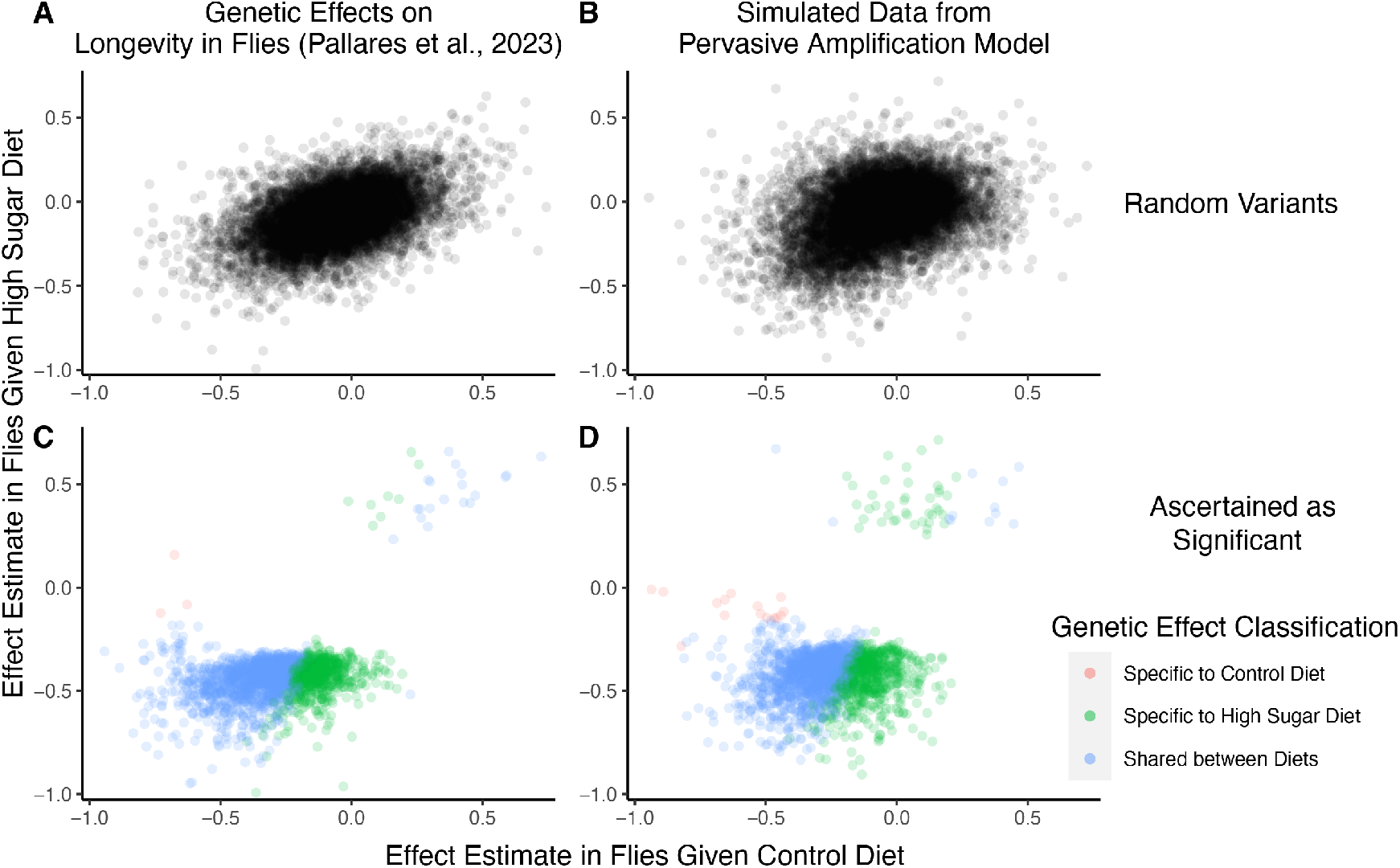
A focus on top hits may be lead to mischaracterization of polygenic GxE. **(A)** Data from an experiment measuring allelic effects on longevity in caged flies given one of two diets, “control” and “high sugar”. Shown are allelic effect estimates under each diet for a random sample of approximately 12K variants. (B) Simulated data where all true allelic effects are exactly 1.4 times larger under a high-sugar diet. The effects are estimated with sampling noise mimicking the Pallares et al. data. **(C)** Allelic effect estimates of variants ascertained as significant and classified as “diet-specific” or “shared” by Pallares et al. **(D)** Simulated effects ascertained as significant and classified using a similar procedure to that applied in (C). While the generative mode of GxE we used in our simulations was not considered by Pallares et al., the simulation results (left panels) closely match the patterns observed in their data (right panels) across all effects (top panels) and as reflected via their classification approach (bottom panels).

To recap, we simulated a mode of GxE that is not considered in Pallares et al. (i.e., pervasive amplification) and that is at odds with their conclusions about evidence for a large discrete class of SNPs with diet-specific effects (i.e., cryptic genetic variation). The close match of our simulation to the empirical results of Pallares et al. therefore illustrates that the characterization of GxE via hypothesis testing and classification at each variant independently may lead to erroneous interpretation when applied to empirical complex trait data as well. In the **Supplementary Materials**, we show that a re-analysis of the Pallares et al. data that is based on estimating the covariance of allelic effects directly is consistent with pervasive amplification as well (**Fig. S4**). In conclusion, the classification of “top hits” alone may not be representative of the extent of GxE nor of the most pervasive modes of GxE.

### The utility of modeling GxE for complex trait prediction

Modeling context dependency of genetic effects may hold the potential for constructing polygenic scores that are more accurate, or improve their portability across contexts [34, 41–45]. Evidence for the utility of GxE models in polygenic score prediction, however, has been underwhelming and GxE models are still rarely applied [9, 10]. A key reason behind this apparent discrepancy is the bias–variance tradeoff for individual variants discussed above. If context-specific effects are similar—a likely possibility for highly polygenic traits with the majority of heritability owing to small causal effects—then additive models will tend to outperform [18, 19, 46, 47]. This is because the unbiasedness of GxE estimation does not make up for the cost of additional estimator variance, resulting from sample stratification by context or the addition of explicit interaction terms [10].

We exemplify the relative importance of variance compared to bias in polygenic scoring using simulations. We continue with the generative model of pervasive amplification as an example. Namely, we simulated a GWAS of a continuous trait with independent effects in 2, 500 variants (50% of variants included in the GWAS). Effects were either the same in two contexts, *A* and *B*, or 1.4 times larger in context *B*. The GWAS is conducted with either a small sample size or a large sample size, conferring low or high statistical power, respectively. We then constructed polygenic scores using 833 variants (corresponding to one-third of the causal variants), which were ascertained as most significantly associated with the trait according to either the additive model (orange and red in **Fig. 5**) in or context-specific hypothesis tests (green and blue in **Fig. 5**).

**Figure 5.**
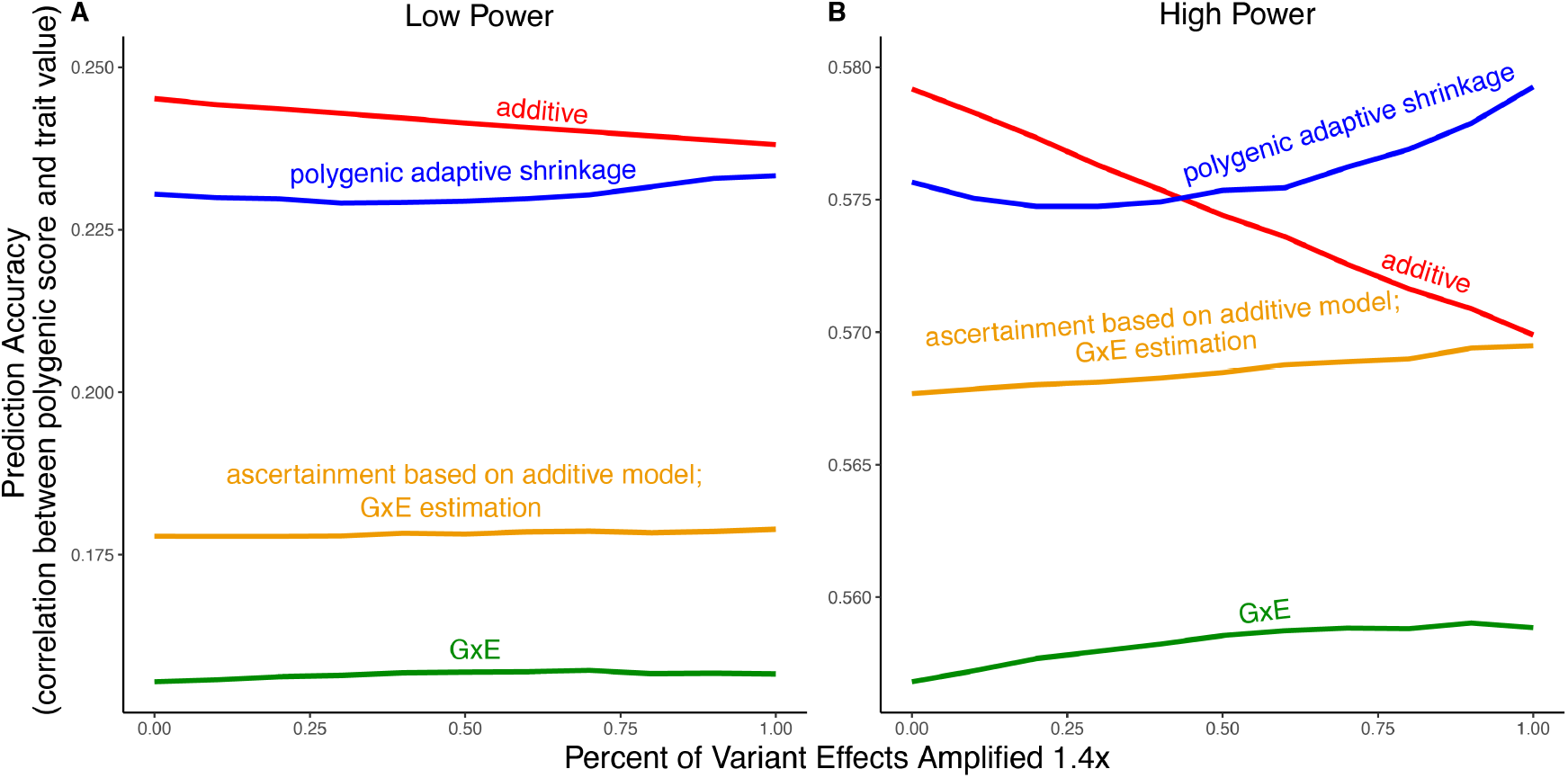
Polygenic score performance for context-dependent prediction models. In each simulation, a GWAS is performed on 5, 000 biallelic variants, half of which have no effect in either context. Of the other half, some percent of the variants (indicated on the x-axis) had effects 1.4 × larger in one of contexts and the remaining SNPs had equal effects in both contexts. The broad sense heritability was set to 0.4 in all simulations. The y-axis shows the average, over 11, 000 simulations, of the out-of-sample Pearson correlation between polygenic score and trait value. **(A)** Results with a GWAS sample size of 1, 000 individuals. **(B)** Results with a GWAS size of 50, 000 individuals.

Even in settings with pervasive GxE, additive polygenic scores (red lines in **Fig. 5**) outperformed context-specific scores (green lines in **Fig. 5**). The advantage of the additive model is manifested in two ways: more accurate estimation, as discussed above, but also better identification of true associations with the trait. We considered the two advantages separately. It is sometimes better to ascertain variants using the lower variance approach and estimate effects using the lower-bias approach. In our simulations, this strategy (orange lines in **Fig. 5)** was preferable to using the GxE model for both ascertainment and estimation (green line). It was not preferable to using the additive model (red line) for both approaches; but it was the preferable strategy under a slightly different parametric regime, corresponding to more GxE (**Fig. S4B**).

Finally, we considered a polygenic GxE approach, as implemented in “multivariate adaptive shrinkage” (*mash*) [29], a method to estimate context-specific effects by leveraging common patterns of effect covariance between contexts observed across the genome. *mash* models the underlying distribution of effects in all contexts as a mixture of zero-centered Multivariate Normal distributions with different covariance structures (as well as the null matrix, to induce additional shrinkage). After estimating this distribution via maximum likelihood, *mash* uses it as a prior to obtain posterior effect estimates for each variant in each context. As a result, posterior effect estimates across contexts regress towards commonly observed patterns of covariance of allelic effects across contexts.

In our simulations, in the presence of substantial amplification, the polygenic adaptive shrinkage approach outperformed all other methods as long as the study was adequately powered (**Fig. 5B**). This is thanks to the unique ability (compared to the three other approaches) to leverage the sharing of signals across variants, including the extent and nature of context dependency. With low power, however, the additive model performed best (**Fig. 5A**). We attribute this to the variance cost associated with the polygenic adaptive shrinkage approach—driven by the estimation of additional parameters for capturing the genome-wide covariance relationships.

## Conclusion

When genetic variants are considered independently, the estimation of their effects in different contexts can be boiled down to a bias-variance tradeoff. For complex traits, we show through example that further considering polygenic patterns of GxE can be key for understanding context-dependent genetic architecture and to aid in prediction. The notion that complex trait analyses should combine observations at top associated loci alongside polygenic trends has gained traction with additive models of trait variation; it may be similarly important in our understanding of context-dependency.

## Acknowledgements

We thank Doc Edge, Marc Feldman, Mark Kirkpatrick, Molly Przeworski, Anil Raj, Elliot Tucker-Drob and members of the Harpak Lab for comments on the manuscript. We thank Peter Andolfatto, Julien Ayroles and Tom Juenger for helpful discussions. All authors were supported by NIH R35GM151108 to A.H. S.P. Smith was also supported by NIH RF1AG073593. This study was conducted using the UK Biobank resource under application 61666, as approved by the University of Texas at Austin institutional review board (protocol 2019-02-0125).

## Methods

### Expressing the Additive Estimator as a Linear Combination of GxE Estimators

In this section, we prove the result of **Eq. 3**, stating that

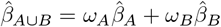

for some non-negative weights *ω*_*A*_ and *ω*_*B*_. To do this, we will need some additional notation. Let 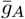 denote the average number of effect alleles in individuals in context *A*, and let 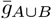 denote the average effect allele count across all individuals. Similarly, let 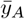 denote the average trait value in context *A*, and let 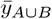 denote the average trait value across all individuals.

As an OLS estimator, the context-specific estimator is defined as

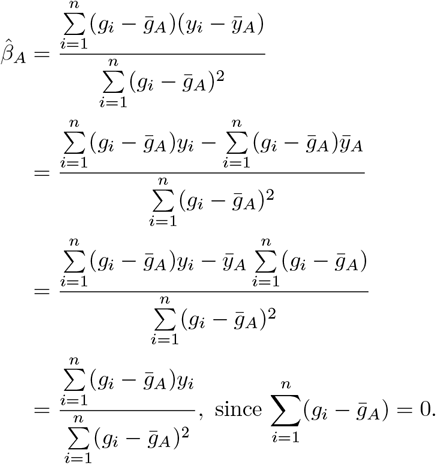

Similarly, the additive estimator can be written as:

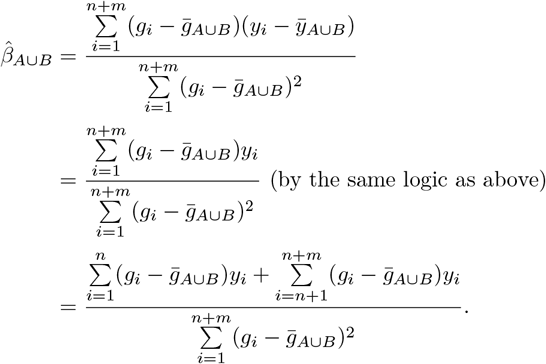

We will show that the weights in **Eq. 3** depend on the effect allele frequency in the two contexts, *f*_*A*_ and *f*_*B*_. We will assume mean-centered traits, such that 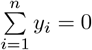 and 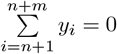. We note that mean-centering is inconsequential for effect estimation. We can then write

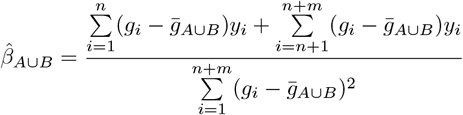

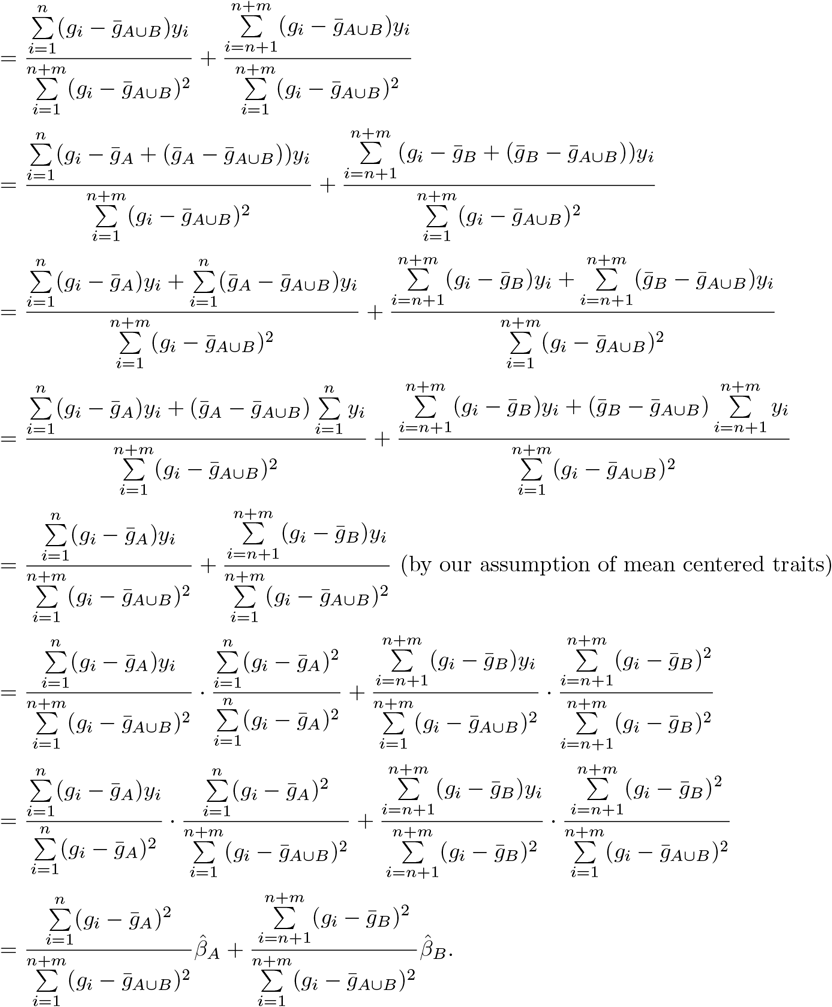

Thus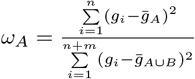 and 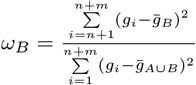 in **Eq. 3**. We note that the numerator of *ω*_*A*_ is *n* times the sample heterozygozity in context A, and the numerator of *ω*_*B*_ is *m* times the sample heterozygozity in context B. Thus, we have shown that

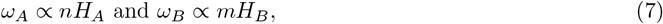

where *H*_*A*_ and *H*_*B*_ are the sample heterozygozities in context A and B, respectively. And, in the special case where *f*_*A*_ = *f*_*B*_, because this implies that the sample heterozygozities will be approximately equal across contexts, we have that

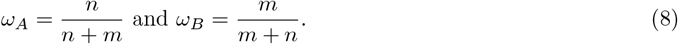

### Linearity of the decision rule

In **Eq. 5**, under the assumption that *V*_*A*_ = *V*_*B*_ and *H*_*A*_ = *H*_*B*_, the decision boundary is expressed as a linear function of |*β*_*A*_ − *β*_*B*_| and 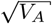 as

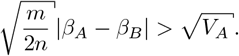

Here, we prove that the linearity of the decision rule holds in the more general case where 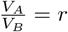 for some fixed value of *r*. **Eq. 5** then follows as a special case of this fact when *r* = 1.

Starting from **Eq. 4**, we prefer the GxE estimator to the additive estimator when estimating *β*_*A*_ if

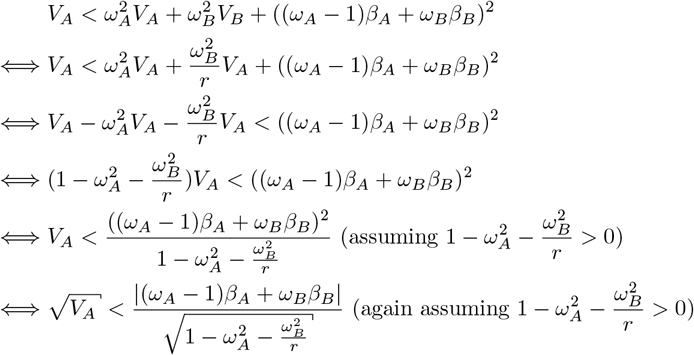

If our assumption that 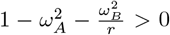 does not hold, we note that the GxE model is always preferable and technically speaking there exists no decision rule between the two models. Now, when heterozygozities (and thus minor allele frequencies) are equal across contexts, then **Eq. 8** implies *ω*_*A*_ + *ω*_*B*_ = 1. Therefore, we may write the decision rule as

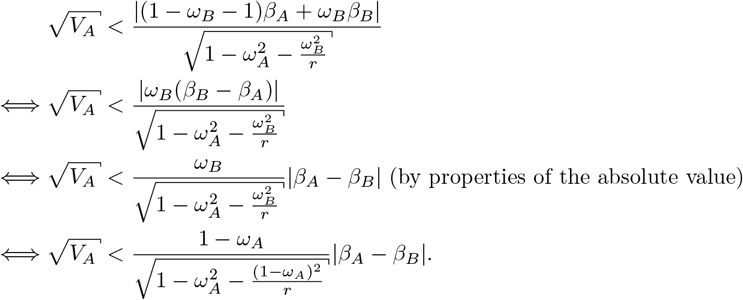

Here, we see that for any fixed *r* the decision rule is linear with a slope determined by *r* (**Fig. 2**). Now, in the special case where *r* = 1, we have

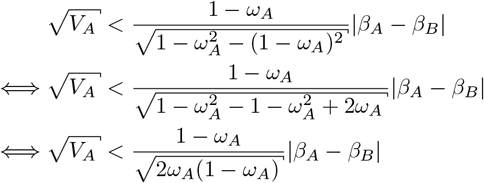

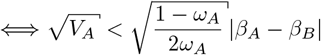

Now, substituting the definitions of *ω*_*A*_ and *ω*_*B*_ in the case of equal minor allele frequencies given in **Eq. 8**, we can write

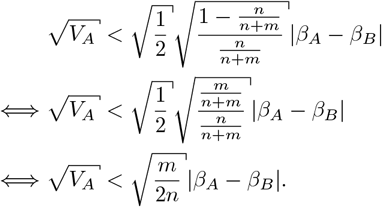

This inequality is instead an equality under the conditions stated in **Eq. 5**. Finally, again using the definition of *ω*_*A*_ and *ω*_*B*_ given in **Eq. 8**, we note that our assumption that 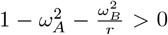 will always hold in the case of equal minor allele frequencies and *r* = 1, as

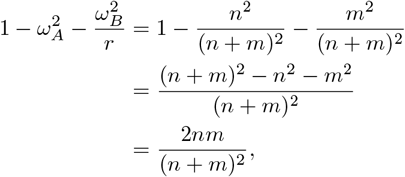

which is strictly positive.

### Re-parameterized decision rule in terms of unitless quantities

In **Eq. 6**, under the assumption that *H*_*A*_ = *H*_*B*_, we re-state the decision rule in terms of the signal-to-noise ratio. Here, we prove this result.

From **Eq. 4**, we have that we should select the GxE model to estimate *β*_*A*_ if and only if

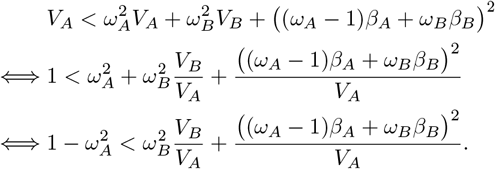

Now, because *H*_*A*_ = *H*_*B*_, we know by **Eq. 8** that *ω*_*A*_ + *ω*_*B*_ = 1. Then, we may write the decision rule as

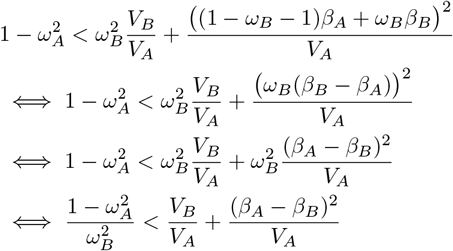

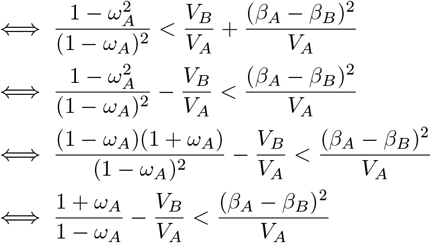

as is stated in **Eq. 6**.

### Simulation of GxDiet effects on longevity in Drosophila

In **Fig. 4**, we compare the effect estimates of Pallares et al. to ones we got in simulations of pervasive amplification. Here, we detail the simulation approach.

We first generated true effects under each diet. For variants *j* = 1, …, 50, 000, we sampled a true effect under the high-sugar diet 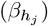 and under the control diet 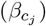.A random 60% of variants were set to have no effect under either diet, with the effects of the remaining 40% of variants sampled as

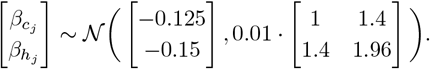

This corresponds to a systematic amplification of 1.4× in the high-sugar compared to the control diet. We selected these parameters based on inspection of the resulting distribution of effects and their correspondence to the Pallares et al. data.

We then simulated the effect estimation. For each variant, the effect estimate was simulated as Normally distributed with mean equal to the true effect and standard deviation equal to a randomly sampled (with replacement) standard error from the effect estimates of Pallares et al. That is, given the simulated values of the true effect estimates 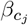 and 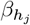,we simulated effect estimates as

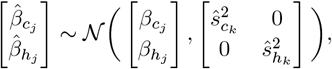

where *k* represents the index of a randomly selected variant from the empirical data of Pallares et al. and 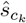 and 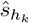 are the corresponding estimated standard errors for the effect estimates in the control and high-sugar groups, respectively. This process yielded vectors of estimated effects in the high-sugar group and control group, 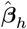 and 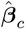,respectively, and vectors of estimated standard errors in the high-sugar group and control group, **ŝ**_*h*_ and **ŝ**_*c*_, respectively. We then performed a Z-test for each variant under each diet, yielding two vectors of p-values, ***p***_*h*_ and ***p***_*c*_, corresponding to the high-sugar and control diets, respectively.

Using these p-values, we followed a similar approach to Pallares et al. to classify the variants (**Fig. 4D**). First, as in Pallares et al., we computed q-values separately for each diet [48], yielding ***q***_*h*_ and ***q***_*c*_, corresponding to the q-values of non-zero effects in the high-sugar and control diets, respectively. Then, we employed the following classification scheme for each variant *j* = 1, …, 50, 000:

1. if 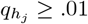 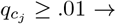;classify as *no effect*.
2. if 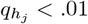 and 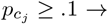;classify as *high-sugar specific effect*.
3. if 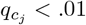 and 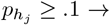;classify as *control specific effect*.
4. if 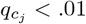 and 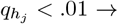;classify as *shared effect*.

We note that p-value and q-value cutoffs used are nominally different than those used in the Pallares et al. study.

### Polygenic Score Simulations

In **Fig. 5**, we show the results of multiple simulations where we compute polygenic scores in each of two contexts under amplification. Here, we detail the generation of data in the simulations and the methods for constructing polygenic scores.

As in **Results and Discussion**, we assumed that we have *n* + *m* observations of a continuous trait, where the first *n* individuals are observed in context *A* and the final *m* are observed in context *B*. For convenience, in this case we assumed *n* = *m*. Now, for variants *j* = 1, …, *p* we generated true effects in contexts *A* and *B* independently from the mixture model

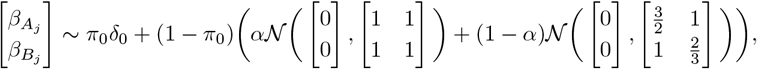

where π_0_ (which we set to 0.5) represents the proportion of SNPs with null effects in both contexts, *α* represents the proportion of non-null SNPs which have exactly equal effects in both contexts, and 1 − *α* is the proportion of non-null SNPs which are generated as perfectly correlated but with 1.5× the standard deviation in context A. Let 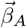 and 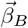 represent the resulting p-vectors of true effects for contexts *A* and *B*, respectively.

Next, we generated genotype counts for each of the *n* + *m* individuals at all *p* variants. Specifically, we independently generated genotypes as

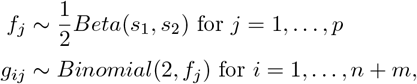

where *f*_*j*_ is the minor allele frequency at variant *j* in the population, *s*_1_ and *s*_2_ are parameters controlling the distribution of minor allele freqeuncies in the population, and *g*_*ij*_ is the observed genotype for individual *i* at variant *j*. Here, we set *s*_1_ = 1 and *s*_2_ = 5. Let ***G***_*A*_ and ***G***_*B*_ represent the generated *n* × *p* matrices of genotypes in contexts A and B, respectively.

Finally, we generated the observed continuous traits for context 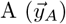 and context 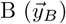 as

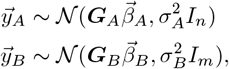

where 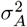 and 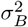 are the observation variances in contexts A and B, respectively, and *I*_*w*_ is the *w*×*w* identity matrix. In our simulations, we set 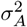 and 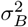 such that the narrow sense heritability is 40% in each context. So that we may later test the accuracy of our polygenic scores, we generated both a training set (consisting of *n* individuals in each context, where *n* = 1, 000 in the low power simulation and *n* = 50, 000 in the high power simulation) for effect estimation and a test set (consisting of 3, 000 individuals in each context) using the above distributions.

**Fig. 5** compares four distinct approaches for constructing polygenic scores, derived from three allelic effect estimation approaches: additive estimation with shrinkage, GxE estimation with shrinkage and *mash*. First, the additive and GxE estimates are derived independently for each variant as described in **Results and Discussion**. Let 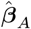 and 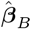 be the p-vectors of GxE estimates of effects in context *A* and *B*, respectively. Similarly, let **ŝ**_*A*_ and **ŝ**_*B*_ be the p-vectors of the standard errors of GxE estimates of effects in context A and B, respectively. Finally, let 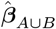 be the p-vector of estimated effects from the additive model and **ŝ**_*A*∪*B*_ be the p-vector of standard errors of estimated effects from the additive model. Using the GxE estimates, we also constructed estimates of the effects in each context using *mash*. Specifically, we ran mash on the *n* × 2 matrices 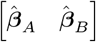 (of effects) and 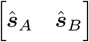 (of standard errors). *mash* then yields 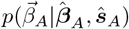 and 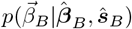,the posterior distributions of the effects in contexts *A* and *B*, respectively.

To construct each polygenic score, we made two choices. First, a choice between the three sets of p-values (or pseudo p-values, see below) for thresholding—we include the 833 (corresponding to one-third of the causal variants) most significant variants in the polygenic score. The second choice was between the three sets of effect estimates to be used as weights in the polygenic score (**Fig. 5**). For instance, when the GxE model was used for ascertainment, we selected the set of variants Ω_*A*_ ⊂ {1, …, *p*} consisting of the variants with the 833 smallest p-values and Ω_*B*_ ⊂ {1, …, *p*} consisting of the variants with the 833 smallest p-values (derived from 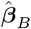 and **ŝ**_*B*_). Then, we predicted trait values (out of sample) by multiplying the effect estimates of our chosen “estimation method” (for *mash* we use the posterior mean) by the effect allele count at each of the selected variants for the individual in question.

## References

1. El-Soda, M., Malosetti, M., Zwaan, B. J., Koornneef, M. & Aarts, M. G. Genotype× environment interaction QTL mapping in plants: lessons from Arabidopsis. Trends in plant science 19, 390–398 (2014).

2. Vieira, C. et al. Genotype-environment interaction for quantitative trait loci affecting life span in Drosophila melanogaster. Genetics 154, 213–227 (2000).

3. Des Marais, D. L., Hernandez, K. M. & Juenger, T. E. Genotype-by-environment interaction and plasticity: exploring genomic responses of plants to the abiotic environment. Annual Review of Ecology, Evolution, and Systematics 44, 5–29 (2013).

4. Smith, E. N. & Kruglyak, L. Gene–environment interaction in yeast gene expression. PLoS biology 6, e83 (2008).

5. Paaby, A. B. & Rockman, M. V. Cryptic genetic variation: evolution’s hidden substrate. Nature Reviews Genetics 15, 247–258 (2014).

6. Munafò, M. R., Zammit, S. & Flint, J. Practitioner Review: A critical perspective on gene–environment interaction models–what impact should they have on clinical perceptions and practice? Journal of child Psychology and Psychiatry 55, 1092–1101 (2014).

7. Kraft, P. & Aschard, H. Finding the missing gene–environment interactions. European Journal of Epidemiology 30, 353–355 (2015).

8. Sella, G. & Barton, N. H. Thinking about the evolution of complex traits in the era of genome-wide association studies. Annual Review of Genomics and Human Genetics 20, 461–493 (2019).

9. Zhu, C. et al. Amplification is the primary mode of gene-by-sex interaction in complex human traits. Cell Genomics, 100297 (2023).

10. Schwaba, T. et al. Comparison of the Multivariate Genetic Architecture of Eight Major Psychiatric Disorders Across Sex. medRxiv, 2023–05 (2023).

11. Elgart, M. et al. Correlations between complex human phenotypes vary by genetic background, gender, and environment. Cell Reports Medicine 3, 100844 (2022).

12. Duncan, L. E. & Keller, M. C. A critical review of the first 10 years of candidate gene-by-environment interaction research in psychiatry. American Journal of Psychiatry 168, 1041–1049 (2011).

13. Gibson, G. & Lacek, K. A. Canalization and robustness in human genetics and disease. Annual Review of Genetics 54, 189–211 (2020).

14. Brown, B. C., Ye, C. J., Price, A. L., Zaitlen, N., Consortium, A. G. E. N. T. 2. D., et al. Transethnic genetic-correlation estimates from summary statistics. The American Journal of Human Genetics 99, 76–88 (2016).

15. Ge, T., Chen, C.-Y., Neale, B. M., Sabuncu, M. R. & Smoller, J. W. Phenome-wide heritability analysis of the UK Biobank. PLoS Genetics 13, e1006711 (2017).

16. Balliu, B. et al. An integrated approach to identify environmental modulators of genetic risk factors for complex traits. The American Journal of Human Genetics 108, 1866–1879 (2021).

17. Veller, C., Przeworski, M. & Coop, G. Causal interpretations of family GWAS in the presence of heterogeneous effects. bioRxiv, 2023–11 (2023).

18. Fisher, R. A. The genetical theory of natural selection (Clarendon Press, 1930).

19. Falconer, D. S. & Mackay, T. F. Introduction to quantitative genetics (Longman, 1996).

20. Yengo, L. et al. A saturated map of common genetic variants associated with human height. Nature 610, 704–712 (2022).

21. Zwick, M. E., Cutler, D. J. & Chakravarti, A. Patterns of genetic variation in Mendelian and complex traits. Annual Review of Genomics and Human Genetics 1, 387–407 (2000).

22. Casella, G. & Berger, R. L. Statistical Inference (Cengage Learning, 2021).

23. Bycroft, C. et al. The UK Biobank resource with deep phenotyping and genomic data. Nature 562, 203–209 (2018).

24. Sinnott-Armstrong, N., Naqvi, S., Rivas, M. & Pritchard, J. K. GWAS of three molecular traits highlights core genes and pathways alongside a highly polygenic background. Elife 10, e58615 (2021).

25. Shi, H., Kichaev, G. & Pasaniuc, B. Contrasting the genetic architecture of 30 complex traits from summary association data. The American Journal of Human Genetics 99, 139–153 (2016).

26. Boyle, E. A., Li, Y. I. & Pritchard, J. K. An expanded view of complex traits: from polygenic to omnigenic. Cell 169, 1177–1186 (2017).

27. Liu, X., Li, Y. I. & Pritchard, J. K. Trans effects on gene expression can drive omnigenic inheritance. Cell 177, 1022–1034 (2019).

28. Wray, N. R., Wijmenga, C., Sullivan, P. F., Yang, J. & Visscher, P. M. Common disease is more complex than implied by the core gene omnigenic model. Cell 173, 1573–1580 (2018).

29. Urbut, S. M., Wang, G., Carbonetto, P. & Stephens, M. Flexible statistical methods for estimating and testing effects in genomic studies with multiple conditions. Nature Genetics 51, 187–195 (2019).

30. Zhang, L. et al. QTL× environment interactions underlie ionome divergence in switchgrass. G3 11, jkab144 (2021).

31. Paaby, A. B. & Gibson, G. Cryptic genetic variation in evolutionary developmental genetics. Biology 5, 28 (2016).

32. Aschard, H. et al. Evidence for large-scale gene-by-smoking interaction effects on pulmonary function. International journal of epidemiology 46, 894–904 (2017).

33. Gibson, G. & Dworkin, I. Uncovering cryptic genetic variation. Nature Reviews Genetics 5, 681–690 (2004).

34. Miao, J. et al. Reimagining gene-environment interaction analysis for human complex traits. bioRxiv, 2022–12 (2022).

35. Spierer, A. N. et al. Natural variation in the regulation of neurodevelopmental genes modifies flight performance in Drosophila. PLoS Genetics 17, e1008887 (2021).

36. Traglia, M., Bout, M. & Weiss, L. A. Sex-heterogeneous SNPs disproportionately influence gene expression and health. PLoS Genetics 18, e1010147 (2022).

37. Bernabeu, E. et al. Sex differences in genetic architecture in the UK Biobank. Nature Genetics 53, 1283–1289 (2021).

38. Wright, K. M. et al. Age and diet shape the genetic architecture of body weight in diversity outbred mice. Elife 11, e64329 (2022).

39. Pallares, L. F. et al. Dietary stress remodels the genetic architecture of lifespan variation in outbred Drosophila. Nature Genetics, 1–7 (2022).

40. Young, A. I., Wauthier, F. & Donnelly, P. Multiple novel gene-by-environment interactions modify the effect of FTO variants on body mass index. Nature communications 7, 12724 (2016).

41. Mostafavi, H. et al. Variable prediction accuracy of polygenic scores within an ancestry group. eLife 9, e48376 (2020).

42. Patel, R. A. et al. Genetic interactions drive heterogeneity in causal variant effect sizes for gene expression and complex traits. The American Journal of Human Genetics 109, 1286–1297 (2022).

43. Turley, P. et al. Multi-trait analysis of genome-wide association summary statistics using MTAG. Nature Genetics 50, 229–237 (2018).

44. Spence, J. P., Sinnott-Armstrong, N., Assimes, T. & Pritchard, J. K. A flexible modeling and inference framework for estimating variant effect sizes from GWAS summary statistics. bioRxiv, 2022–04 (2022).

45. Wang, J. Y. et al. Three Open Questions in Polygenic Score Portability. bioRxiv (2024).

46. Hill, W. G., Goddard, M. E. & Visscher, P. M. Data and theory point to mainly additive genetic variance for complex traits. PLoS Genetics 4, e1000008 (2008).

47. Young, A. I. Solving the missing heritability problem. PLoS Genetics 15, e1008222 (2019).

48. Storey, J. D. The positive false discovery rate: a Bayesian interpretation and the q-value. The Annals of Statistics 31, 2013–2035 (2003).

