## Supplementary Materials for "Tradeoffs in Modeling Context Dependency in Complex Trait Genetics"

#### Contents

**1 Validating and Applying the Decision Rule to Real Data 2**

**2 Equivalence of Regression Coefficient Estimates for Explicit Interaction Term and Stratified Model 2**

**3 Continuous Context Variable 4**

**4 Gene-Context Correlation 5**

**5 Classification-Based Inference in Pallares et al., 2022 6**

**6 Reanalysis of GxE in the Pallares et al. Experiment 7**

### 1 Validating and Applying the Decision Rule to Real Data

To validate the decision rule of **Eqs. 4-6** of the main text, we considered the example of sex as a context for physiological traits using UK Biobank data [1]. For each of 27 physiological quantitative traits, we first split the sample by sex chromosome karyotypes into XX individuals (females) and XY individuals (famles). For each sex, we randomly split the sample into a training set with 80% of the individuals and a test set with 20% of the individuals. We estimated within-sex GWAS in the training and test sets separately. We denote by $\hat{\beta}^{(t)}_{Z_i}$ the marginal effect estimate of the $i^{th}$ SNP in sex $Z$ in set $t$, and by $\hat{s}^{(t)}_{Z_i}$ the corresponding standard error. We denote by $\hat{\beta}^{(train)}_{M\cup F_i}$ the estimated effect of the $i^{th}$ SNP in the union of training sets (with both sexes included). In our validation procedure, we treated $\hat{\beta}^{(test)}_{M_i}$ as the "ground-truth," i.e., as the true effect. While $\hat{\beta}^{(test)}_{M_i}$ is of course a very noisy estimate of $\beta_{M_i}$, it is an unbiased estimate, so we expect that estimates that are closer to $\hat{\beta}^{(test)}_{M_i}$ in the aggregate will also be closer to $\beta_{M_i}$. Using this assumption, we define two empirical quantities,

$$
\begin{aligned}
error(\hat{\beta}^{(train)}_{M_i}) &= (\hat{\beta}^{(train)}_{M_i} - \hat{\beta}^{(test)}_{M_i})^2 \\
error(\hat{\beta}^{(train)}_{M\cup F_i}) &= (\hat{\beta}^{(train)}_{M\cup F_i} - \hat{\beta}^{(test)}_{M_i})^2.
\end{aligned}
$$

Given $\sqrt{V_{M_i}}$ and $|\beta_{M_i} - \beta_{F_i}|$, **Eq. 4** allows us to calculate the expected difference between $error(\hat{\beta}^{(train)}_{M_i})$ and $error(\hat{\beta}^{(train)}_{M\cup F_i})$. However, we do not observe $\sqrt{V_{M_i}}$ or $|\beta_{M_i} - \beta_{F_i}|$. Instead, we estimated these quantities and examined the agreement of the estimates in the two sets. We estimated $\sqrt{V_{M_i}}$ as $\hat{s}^{(train)}_{M_i}$. Estimating $|\beta_{M_i} - \beta_{F_i}|$ is more difficult, as the intuitive $|\hat{\beta}^{(train)}_{M_i} - \hat{\beta}^{(train)}_{F_i}|$ is an upwardly biased estimator. Instead, we used an empirical shrinkage estimator (*ash* [2]) to estimate $|\beta_{M_i} - \beta_{F_i}|$. In essence, *ash* takes in a vector of effect estimates $\hat{\boldsymbol{\beta}}$ and a vector of corresponding standard errors $\hat{\boldsymbol{s}}$, estimates a prior distribution of the true effects $p(\boldsymbol{\beta})$, and then uses this prior to obtain a posterior distribution of true effects.

Specifically, for a large random sample of sites, $i = 1, \dots, p$, we ran *ash* on the vectors

$$
\begin{aligned}
\hat{\boldsymbol{\beta}} &= \left(\hat{\beta}^{(train)}_{M_1} - \hat{\beta}^{(train)}_{F_1}, \dots, \hat{\beta}^{(train)}_{M_p} - \hat{\beta}^{(train)}_{F_p}\right) \\
\hat{\boldsymbol{s}} &= \left(\sqrt{(\hat{s}^{(train)}_{M_1})^2 + (\hat{s}^{(train)}_{F_1})^2}, \dots, \sqrt{(\hat{s}^{(train)}_{M_p})^2 + (\hat{s}^{(train)}_{F_p})^2}\right).
\end{aligned}
$$

and took the absolute value of *ash*'s output posterior mean estimates of $\beta_{M_i} - \beta_{F_i}$ as our estimate of $|\beta_{M_i} - \beta_{F_i}|$ for each site. With our estimates of $|\beta_{M_i} - \beta_{F_i}|$ and $\sqrt{V_{M_i}}$ in hand, we calculated the expected difference in squared errors using **Eq. 4** (x-axis in **Fig. 3A**) and compared it with the actual difference between $error(\hat{\beta}^{(train)}_{M_i})$ and $error(\hat{\beta}^{(train)}_{M\cup F_i})$ (**Fig. S1**).

For **Fig. 3A,B**, we used the same procedure as above to estimate the x and y axes, except we did not employ any data splitting.

### 2 Equivalence of Regression Coefficient Estimates for Explicit Interaction Term and Stratified Model

In the main text, we discuss the estimation of context-dependent effects through the estimation of two effects in subsamples stratified by context. The generative model defined in **Eq. 1** is

$$
y_i \sim \begin{cases} \mathcal{N}(\alpha_A + \beta_A g_i, \sigma_A^2) & \text{if } i \in \{1, \dots, n\} \\ \mathcal{N}(\alpha_B + \beta_B g_i, \sigma_B^2) & \text{if } i \in \{n+1, \dots, n+m\}. \end{cases} \tag{S1}
$$

where $\beta_A$ and $\beta_B$ are fixed, context-specific effects of a reference allele at a biallelic, autosomal variant $i$, $g_i \in \{0, 1, 2\}$ is the observed reference allele count. $\alpha_A$ and $\alpha_B$ are the context-specific intercepts, corresponding to the mean trait value for individuals with zero reference alleles in context $A$ and $B$, respectively. $\sigma_A^2$ and $\sigma_B^2$ are context-specific observation variances. Another common way to define a generative GxE model is with a linear genotype-by-context interaction term, namely,

$$y_i \sim \mathcal{N}(\alpha + \beta_G g_i + \beta_E e_i + \beta_{GxE} g_i e_i, \sigma^2) \quad \text{independently for } i = 1, \ldots, n+m, \tag{S2}$$

where $\alpha$ is the mean trait value for individuals with zero reference alleles in context $A$, $\beta_G$ is the genetic effect, $\beta_E$ is the contextual effect, $\beta_{GxE}$ in the genotype-context interaction term, $e_i$ is the contextual covariate for individual $i$, defined to be 0 if the individual is in context $A$ and 1 if the individual is in context $B$, and $\sigma^2$ is the homoskedastic noise term.

Here, we show that least squares estimation of the two models above context-specific effect estimates. The ordinary least squares (OLS) estimate of the intercept and context-specific allelic effects under the model described in **Eq. S1** are

$$\hat{\alpha}_A, \hat{\beta}_A = \underset{\alpha_A, \beta_A}{\text{argmin}} \sum_{i=1}^{n} \big(y_i - (\alpha_A + \beta_A g_i)\big)^2 \tag{S3}$$

$$\hat{\alpha}_B, \hat{\beta}_B = \underset{\alpha_B, \beta_B}{\text{argmin}} \sum_{i=n+1}^{n+m} \big(y_i - (\alpha_B + \beta_B g_i)\big)^2. \tag{S4}$$

The OLS estimates under the model described in **Eq. S2** are

$$\hat{\alpha}, \hat{\beta}_G, \hat{\beta}_E, \hat{\beta}_{GxE} = \underset{\alpha, \beta_G, \beta_E, \beta_{GxE}}{\text{argmin}} \sum_{i=1}^{n+m} \big(y_i - (\alpha + \beta_G g_i + \beta_E e_i + \beta_{GxE} g_i e_i)\big)^2. \tag{S5}$$

Since $e_i = 1$ if individual $i$ is in context $B$ and 0 otherwise, we can re-write **Eq. S5** as

$$\begin{aligned}
\hat{\alpha}, \hat{\beta}_G, \hat{\beta}_E, \hat{\beta}_{GxE} &= \underset{\alpha, \beta_G, \beta_E, \beta_{GxE}}{\text{argmin}} \left( \sum_{i=1}^{n} \big(y_i - (\alpha + \beta_G g_i)\big)^2 + \sum_{i=n+1}^{n+m} \big(y_i - (\alpha + \beta_G g_i + \beta_E + \beta_{GxE} g_i)\big)^2 \right) \\
&= \underset{\alpha, \beta_G, \beta_E, \beta_{GxE}}{\text{argmin}} \left( \sum_{i=1}^{n} \big(y_i - (\alpha + \beta_G g_i)\big)^2 + \sum_{i=n+1}^{n+m} \big(y_i - ((\alpha + \beta_E) + (\beta_G + \beta_{GxE}) g_i)\big)^2 \right).
\end{aligned}$$

By **Eq. S3** we have that $\sum_{i=1}^{n} \big(y_i - (\alpha + \beta_G g_i)\big)^2$ is minimized by setting $\alpha = \hat{\alpha}_A$ and $\beta_G = \hat{\beta}_A$. In turn, by **Eq. S4** we have that $\sum_{i=n+1}^{n+m} \big(y_i - ((\alpha + \beta_E) + (\beta_G + \beta_{GxE}) g_i)\big)^2$ is minimized by setting $\alpha + \beta_E = \hat{\alpha}_B$ and setting $\beta_G + \beta_{GxE} = \hat{\beta}_B$. Thus, both terms are simultaneously minimized by setting

$$\begin{aligned}
\alpha &= \hat{\alpha}_A \\
\beta_G &= \hat{\beta}_A \\
\beta_E &= \hat{\alpha}_B - \hat{\alpha}_A \\
\beta_{GxE} &= \hat{\beta}_B - \hat{\beta}_A.
\end{aligned}$$

Then, under the explicit interaction model, for an individual in context $A$ the estimated intercept is $\hat{\alpha}_A$ and the estimated genetic effect is $\hat{\beta}_A$. And, for an individual in context $B$, the estimated intercept is $\hat{\alpha}_A + \hat{\alpha}_B - \hat{\alpha}_A = \hat{\alpha}_B$ and the estimated genetic effect is $\hat{\beta}_A + \hat{\beta}_B - \hat{\beta}_A = \hat{\beta}_B$. Thus, conditional on the context, both the explicit interaction model and the stratified model provide the same effect and intercept estimates.

We note that there is still an important difference between the stratified and explicit interaction models, as the stratified model allows for heteroskedasticity between contexts where the explicit interaction model does not. We also note that the proof of equivalence relies on the context variable being binary.

### 3 Continuous Context Variable

Here, we extend our analysis of single site estimation in a binary context to the case of a continuous context variable. Specifically, for individuals indexed by $i = 1, \ldots, n$, we consider the generative model

$$y_i \sim \mathcal{N}(\beta_0 + \beta_G g_i + \beta_E e_i + \beta_{G\times E} g_i e_i, \sigma^2), \tag{S6}$$

where $y_i \in \mathbb{R}$ is the continuous trait value for individual $i$, $g_i \in \{0, 1, 2\}$ is the number of reference alleles carried by a diploid individual $i$ at a focal biallelic variant of interest, $e_i \in \mathbb{R}$ is the continuous context for individual $i$, $\beta_0$ is the the mean trait value for individuals with the context variable value of zero and zero reference alleles, $\beta_G$ is the main (additive) allelic effect, $\beta_E$ is the effect of the context, $\beta_{G\times E}$ is the main interaction effect, and $\sigma^2$ is the residual variance. For simplicity, we will assume $\beta_0 = 0$.

For a binary context, we considered the differnce in mean squared error (MSE) of the "additive model" and "GxE model" in estimating the context-specific genetic effect. Analogously, for a continuous context we will consider the difference in MSE in estimating the total genetic effect conditional on a particular value of the context variable.

To perform this analysis, we will consider the estimation of two models. The first model, which we will call the "GxE" model, is described in **Eq. (S6)** above (where we omit the intercept because we assume it is 0). For convenience, we will use the following matrix notation

$$y = \begin{bmatrix} y_1 \\ \vdots \\ y_n \end{bmatrix}, X = \begin{bmatrix} g_1 & e_1 & g_1 e_1 \\ \vdots & \vdots & \vdots \\ g_n & e_n & g_n e_n \end{bmatrix}, \beta = \begin{bmatrix} \beta_G \\ \beta_E \\ \beta_{G\times E} \end{bmatrix}.$$

Then, we consider the standard least squares estimate of $\hat{\beta} = (X^T X)^{-1} X^T y$, for which

$$\hat{\beta} \sim \mathcal{N}(\beta, \sigma^2 (X^T X)^{-1}). \tag{S7}$$

We will assume that the vector of genotypes $\vec{g}$ and the vector of context covariates $\vec{e}$ are orthogonal in our sample and are both mean centered. (In **Section 3** below, we discuss the case of gene-context correlation, where these assumptions are not met.) Then, the matrix $X^T X$ will be approximately diagonal. Under this approximation, we can write **Eq. (S7)** as

$$\hat{\beta} \sim \mathcal{N}(\beta, \Sigma),$$

where

$$\Sigma \approx \sigma^2 \text{diag}\left( \frac{1}{\sum_{i=1}^n g_i^2}, \frac{1}{\sum_{i=1}^n e_i^2}, \frac{1}{\sum_{i=1}^n g_i^2 e_i^2} \right). \tag{S8}$$

Now, we are interested in the error of the GxE model in estimating the genetic effect conditional on a particular value of the context $e$. By **Eq. (S6)**, we define the true genetic effect conditional on a particular value of the context $e$ as $\beta_G + \beta_{G\times E} e$. Thus, to evaluate the error of the GxE model, the conditional expected error is

$$E[((\hat{\beta}_G + \hat{\beta}_{G\times E} e) - (\beta_G + \beta_{G\times E} e))^2 | e]. \tag{S9}$$

By the definition of conditional variance, we can write **Eq. (S9)** as

$$\begin{aligned}
&Var[(\beta_G + \beta_{G\times E} e) - (\hat{\beta}_G + \hat{\beta}_{G\times E} e)|e] + (E[(\beta_G + \beta_{G\times E} e) - (\hat{\beta}_G + \hat{\beta}_{G\times E} e)|e])^2 \\
&= Var[(\beta_G + \beta_{G\times E} e) - (\hat{\beta}_G + \hat{\beta}_{G\times E} e)|e] \quad \text{(since } \hat{\beta}_G \text{ and } \hat{\beta}_{G\times E} \text{ are unbiased estimates)} \\
&= Var[\hat{\beta}_G + \hat{\beta}_{G\times E} e|e] \quad \text{(since } \beta_G \text{ and } \beta_{G\times E} \text{ are fixed parameters)} \\
&\approx \frac{\sigma^2}{\sum_{i=1}^n g_i^2} + \frac{\sigma^2}{\sum_{i=1}^n g_i^2 e_i^2} e^2 \quad \text{(by } \textbf{Eq. (S8)}\text{)}.
\end{aligned}$$

We will also consider the estimation error associated with the "additive model," which in this case estimates the model described in **Eq. (S6)** assuming that $\beta_{G\times E} = 0$. In effect, this assumes that the genetic effect is equal across all values of the context. Because $X^T X$ is approximately diagonal matrix, removing the $G \times E$ covariate and regression coefficient has a negligible effect on both the least squares estimates of $\beta_G$ and its sampling distribution. Thus, we can approximate the error of the additive model by calculating the conditional expectation

$$E[\left((\beta_G + \beta_{G\times E} e) - \hat{\beta}_G\right)^2 | e] \tag{S10}$$

under the model in **Eq. (S6)**. Again, by the definition of the conditional expectation, we can write **Eq. (S10)** as

$$\begin{aligned} &Var[(\beta_G + \beta_{G\times E} e) - \hat{\beta}_G | e] + (E[(\beta_G + \beta_{G\times E} e) - \hat{\beta}_G | e])^2 \\ &= \frac{\sigma^2}{\sum_{i=1}^n g_i^2} + \beta_{G\times E}^2 e^2. \end{aligned}$$

Thus, conditional on a particular value of $e$, we we prefer the $G \times E$ estimator for estimating the genetic effect if and only if

$$\frac{\sigma^2}{\sum_{i=1}^n g_i^2} + \frac{\sigma^2}{\sum_{i=1}^n g_i^2 e_i^2} e^2 < \frac{\sigma^2}{\sum_{i=1}^n g_i^2} + \beta_{G\times E}^2 e^2 \tag{S11}$$

$$\iff \frac{\sigma^2}{\sum_{i=1}^n g_i^2 e_i^2} e^2 - \beta_{G\times E}^2 e^2 < 0 \tag{S12}$$

$$\iff \frac{\sigma^2}{\sum_{i=1}^n g_i^2 e_i^2} < \beta_{G\times E}^2 \tag{S13}$$

This decision rule has a very intuitive interpretation: the $G \times E$ model is preferable when the estimation variance of the interaction term is smaller than the squared interaction effect size. While the exact difference in MSE between the additive and $G \times E$ models depends on the value of the context variable (see **Eq. (S11)**), the decision rule (**Eq. (S13)**) does not.

### 4 Gene-Context Correlation

The simple decision rule given in **Eq. (S13)** was derived using the assumption that the observed genotype vector and the observed context vector were orthogonal. In statistical terms, one may expect this condition to approximately hold when sampling from a population where $r(G, E)$, the correlation between genotypes and the context, is 0. In general, this assumption may not be reasonable.

When this assumption is violated, the design matrix $X$ will no longer form a diagonal matrix when left-multiplied by its transpose. This results in both (a) larger standard errors for each of the marginal coefficients and (b) non-zero sampling covariance between the coefficients. In addition, $|r(G, E)| > 0$ may result in omitted variable bias in the additive model. As a result, the decision rule of **Eq. (S13)** will not hold exactly under conditions of $|r(G, E)| > 0$, and in fact may be far from correct when $|r(G, E)|$ is large.

In general, creating a closed-form decision rule to choose between the GxE and additive models in this scenario is not possible. However, in this section we use a simulation study to gain intuition about the change in bias-variance tradeoff in the presence of gene-context correlation.

In our simulations, for a group of $n$ individuals, we generate genotypes as

$$g_1, \ldots, g_n \overset{iid}{\sim} Binomial(2, p).$$

Then, for some constant $c$, we generate context covariates as

$$e_i \sim \mathcal{N}(c \cdot g_i, \sigma_E^2)$$

independently for $i = 1, \ldots, n$. Finally, we generate trait values $y_1, \ldots, y_n$ using **Eq. (S6)** for particular values of the remaining parameters.

To explore how the bias-variance tradeoff changes for different values of $r(G, E)$, we ran a set of simulations with different values of $c$ in the range of $-3.5$ to $3.5$. For each of these simulations, we generated a dataset of $n = 250$ individuals with $\sigma_E^2 = 1$, $p = 0.1$, $\beta_0 = 0$, $\beta_G = 0.25$, $\beta_E = 0$, $\beta_{G\times E} = -.125$, and $\sigma_y^2 = 7$.

The results of the simulation are shown in **Supplementary Fig. S6**. Non-zero $r(G, E)$ had a number of consequences. First, as the magnitude of $r(G, E)$ increases, the magnitude of the bias of the genetic effect estimate of the additive model also increases (**Supplementary Fig. S6A**). In our simulation, we set the genetic effect and the interaction effect to have opposite signs. When $r(G, E)$ is positive, it pulls the genetic effect estimate towards the interaction effect. In contrast, when $r(G, E)$ is negative, it pulls the genetic effect estimate away from the interaction effect. Since the $G \times E$ model includes all causal variables, it is unbiased. Second, as the magnitude of $r(G, E)$ increases, the variance of the genetic effect estimates of both the additive and $G \times E$ models increase (**Supplementary Fig. S6B**). This is because when variables in a regression are (positively or negatively) correlated, it is more difficult to distinguish between their effects. The $G \times E$ model appears to be affected more strongly here, which likely results from the inclusion of an additional interaction term that is highly correlated with both the genotype covariate and the context covariate. Third, the interaction term in the $G \times E$ model remains unbiased under $r(G, E)$, and increases in variance with larger magnitude of $r(G, E)$ (**Supplementary Fig. S6 C,D**).

### 5 Classification-Based Inference in Pallares et al., 2022

In the main text, we discuss the characterization of GxE by Pallares et al., based on significance testing under each of the diets. The authors classified variants according to whether or not their associations with survivorship were significant under each diet as follows:

1. significant under neither diet → classify as *no effect.*
2. significant when fed the high-sugar diet, but not when fed the control diet → classify as *high-sugar specific effect.*
3. significant when fed the control diet, but not when fed the high-sugar diet → classify as *control specific effect.*
4. significant under both diets → classify as *shared effect.*

Approximately 31% were high-sugar specific, while the remaining 69% of the variants were shared. Fewer than 1% were labelled as having control specific effects. The authors concluded that high-sugar specific effects on longevity are pervasive, compatible with the hypothesis of widespread cryptic genetic variation for longevity.

This "top hits" approach places an emphasis on the context(s) in which trait associations are statistically significant, rather than on estimating how the context-specific allelic effects covary. In addition, this particular classification system also does not cover all possible ways in which context-specific effects may differ. A key example is the case where true effects are concordant in sign but differ in magnitude. The strong positive covariance of estimated effects observed genome-wide (**Fig. 4A**) suggests this case merits consideration. Such variants may fall into each of the existing categories, depending on the magnitude of effects and statistical power in each of the contexts. Three of the four possible classifications are clearly wrong, but what about the "shared" category?

The class "shared" may be interpreted as suggesting lack of context dependency (Fig. 1C in [3]). However, it will tend to include variants having strong effects under both diets, regardless of whether or not the diet-specific effects are similar. As Pallares et al. also note, there are marked differences in the magnitude of diet-specific estimated effects of variants in the "shared" category. Amongst the approximately $1,500$

variants labeled as shared, the estimated effect under the high-sugar diet is on average about 1.3× that of the estimated effect in the control diet.

Notably, the classification as diet-specific does not imply that a variant has an unusually large effect under this diet. On average, variants classified as shared actually have a slightly larger estimated effect under the high-sugar diet than variants classified as high-sugar specific (t-test p=$1.6 \times 10^{-5}$). Thus, instead of suggesting little-to-no effect under the control diet and a large effect under the high-sugar diet (as predicted by the hypothesis that cryptic genetic variation is pervasive), the classification as high-sugar specific may commonly just point us to intermediate size effects—large enough to be significant in the systematically-larger effects context (where power is higher) yet too small to be significant in the systematically-smaller effects context (where power is lower).

### 6 Reanalysis of GxE in the Pallares et al. Experiment

In the main text, we show that the classification of significantly-associated variants in Pallares et al. is consistent with pervasive amplification, despite the fact that this mode of covariance was not one of the generative modes considered by the authors. Here, we re-estimate the modes of covariance of genetic effects under the two diets, using an approach adapted from the one we have previously used [4]. Specifically, we used the framework of "multivariate adaptive shrinkage" (*mash*) to model the covariance of effects between the high-sugar and control diets across all variants.

We fit a Multivariate Normal mixture model to the summary statistics of Pallares et al. [5]. In short, *mash* takes in a set of multidimensional effect estimates and standard errors and estimates the true underlying distribution of effects as a mixture of zero-centered multivariate normal distributions and a point mass at 0. Each mixture component in the model must be pre-specified before the fitting procedure. Following Zhu et al. [4], we pre-specified a dense grid of covariance matrices to be input to mash. In particular, each covariance matrix is of the form

$$\begin{bmatrix} \sigma_c^2 & \rho\sigma_{hs}^2\sigma_c^2 \\ \rho\sigma_{hs}^2\sigma_c^2 & \sigma_{hs}^2 \end{bmatrix},$$

where $\sigma_c^2$ is the variance of the true effects in the control group, $\sigma_{hs}^2$ is the variance of the true effects in the high sugar group, and $\rho$ is the correlation between the effects in the high sugar and control groups.

To specify the set of covariance matrices, we formed a grid encompassing possible values of the correlation of allelic effects across diets and ratio of variances under each of the diets. Specifically, we varied $\rho$ in the set of values $(-1, -\frac{3}{4}, -\frac{1}{2}, -\frac{1}{4}, 0, \frac{1}{4}, \frac{1}{2}, \frac{3}{4}, 1)$, and we varied the ratio of the variances, $\frac{\sigma_c^2}{\sigma_{hs}^2}$ on a logarithmic scale in the range 0.5 and 1.5. Taking the Cartesian product of this sets yielded a grid of covariance matrices.

Because the number of pre-specified covariance matrices was large, and to avoid overfitting, we used a forward step-wise selection procedure [6]. We first started with a model with only the Null covariance matrix (i.e., no effect under either diet), and then added one covariance matrix to the mixture in a greedy manner in each step, by searching over the space of all covariance matrices for a matrix that maximally improves the likelihood of the mixture model. In each step, we either decide to include the new covariance matrix in the model and move to the next step, or instead stop the procedure and not include the new matrix, if the improvement in likeligood compared to the previous step was below a pre-specified threshold. This threshold was determined by conducting a level $\alpha = .05$ likelihood-ratio test.

To mitigate possible linkage disequilibrium (LD) among the input SNPs, we performed the procedure described above on a subset roughly $12K$ out of the roughly $270K$ variants in the data. These SNPs each came as a random sample from $12K$ roughly evenly spaced chromosomal blocks (in terms of physical distance).

Based on the output of this procedure—the mixture weights in the model ultimately chosen—92% of variants have non-zero effects under both diets but larger effects under the high-sugar diet (**Fig. S4**). This suggests that instead of affecting just a small subset of variants, a high-sugar diet amplifies the effects of the vast majority of variants on lifespan. Moreover, *mash* assigned zero weight to covariance matrices where an effect is non-zero in one context but zero in another.

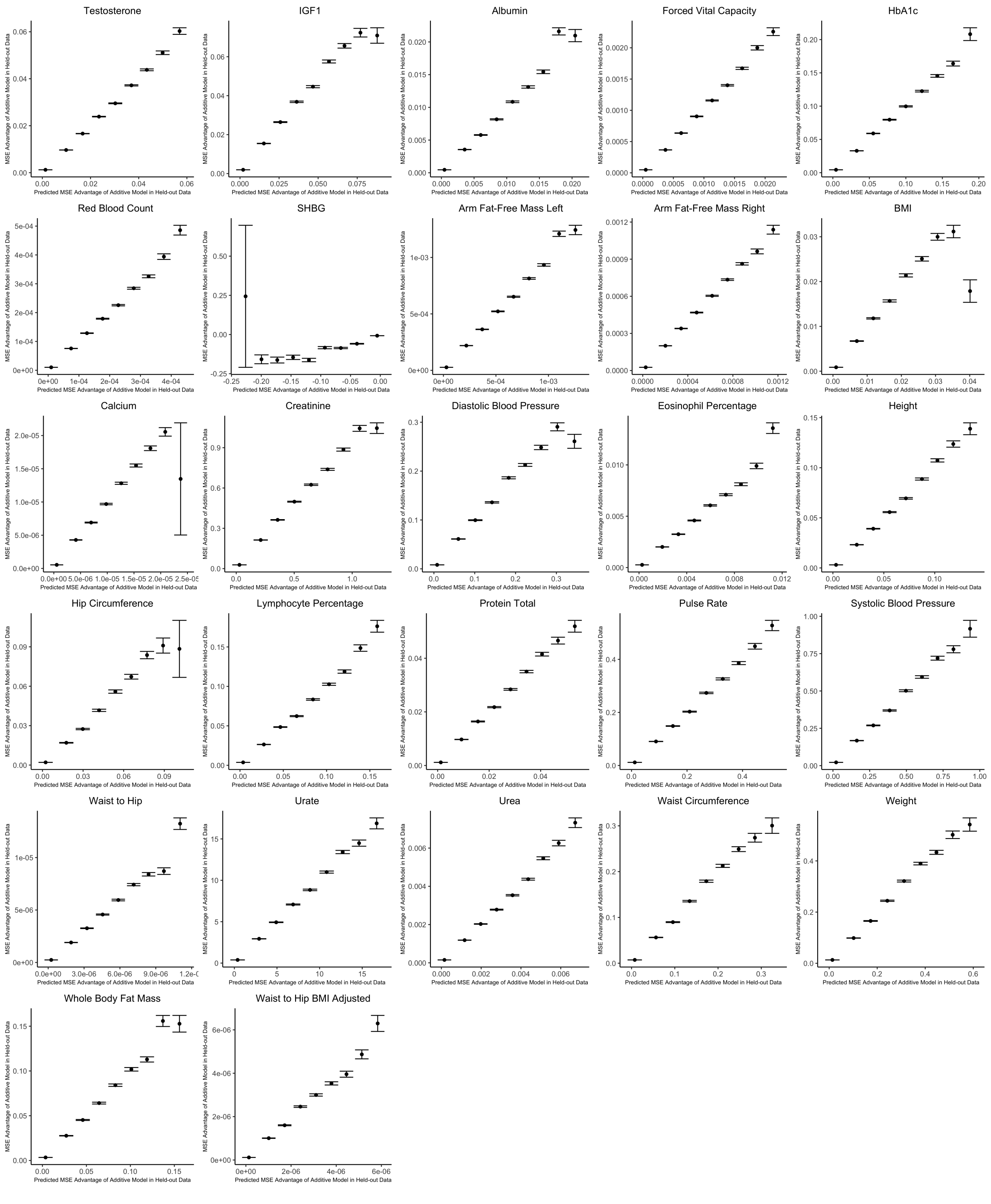

*Supplementary Figure S1: Out of sample validation of decision rule for sex-stratified GWAS for 27 complex traits. For each trait, we use the procedure specified in the section* ***Validation procedure for the decision rule****. The x-axis shows the expected mean squared error (MSE) advantage of the additive estimator in predicting the male effect based on the decision rule, where the y-axis shows the actual MSE advantage of the additive model in predicting the male effect calculated using an independent sample. Values on the x-axis are grouped into 9 evenly spaced bins, and the y-axis (along with corresponding standard errors) shows bin averages. For ease of viewing, data above the* $99^{th}$ *percentile and below the* $1^{st}$ *percentile on the x-axis are removed.*

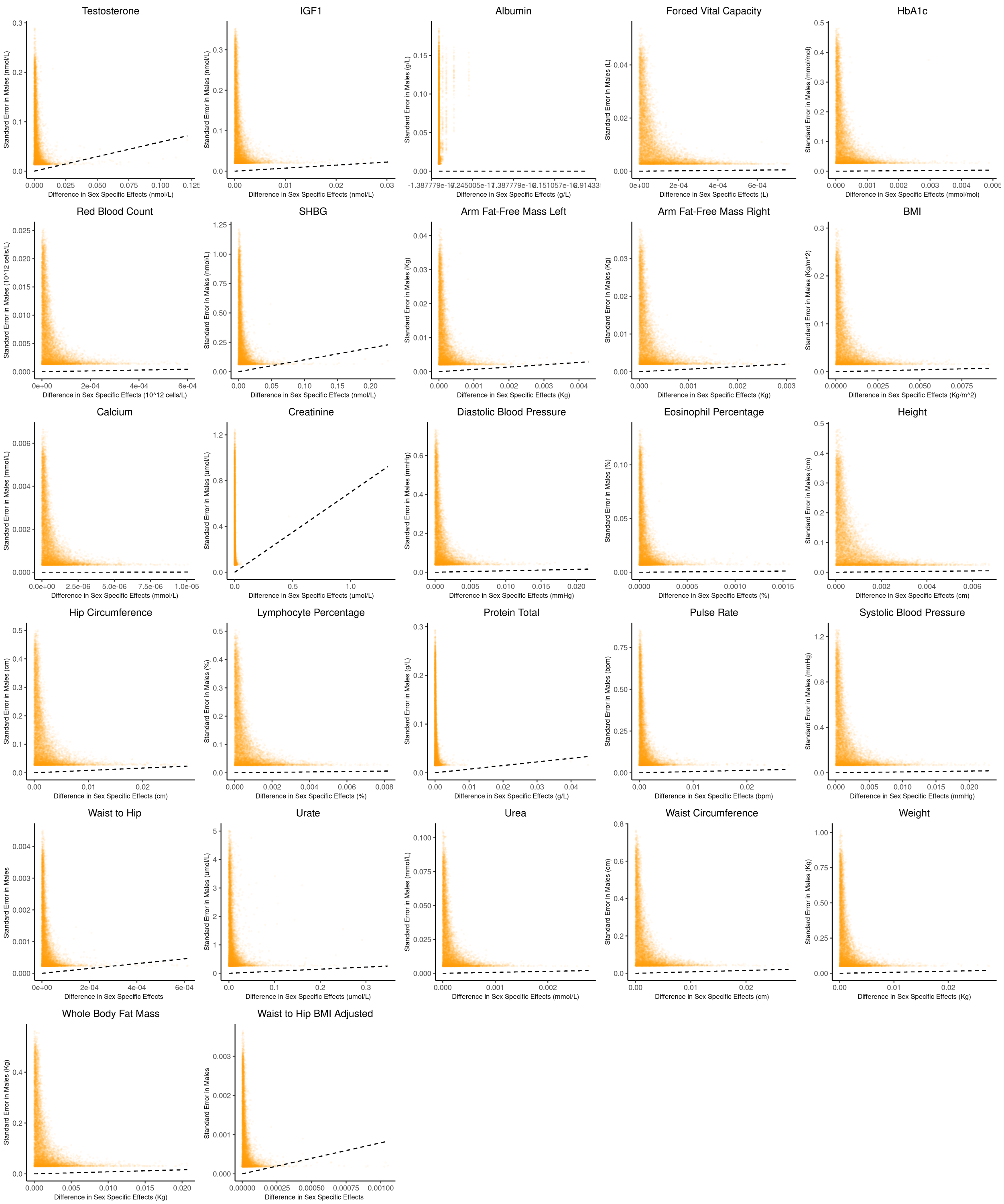

*Supplementary Figure S2: Bias-variance tradeoff at random variants in a sex-stratified GWAS. This figure extends Fig. 3A of the main text to all physiological traits analyzed. The x-axis shows the estimated absolute difference between the effect of variants in males and females. The y-axis shows the measured standard error for each variant in males, the focal context here. The dashed line shows the decision boundary for effect estimation in males. The difference in MSE between estimation methods increases linearly with distance from the dashed line, as in* ***Fig. 2****. If a variant falls above (below) the line, the additive (GxE) estimator has a lower MSE.*

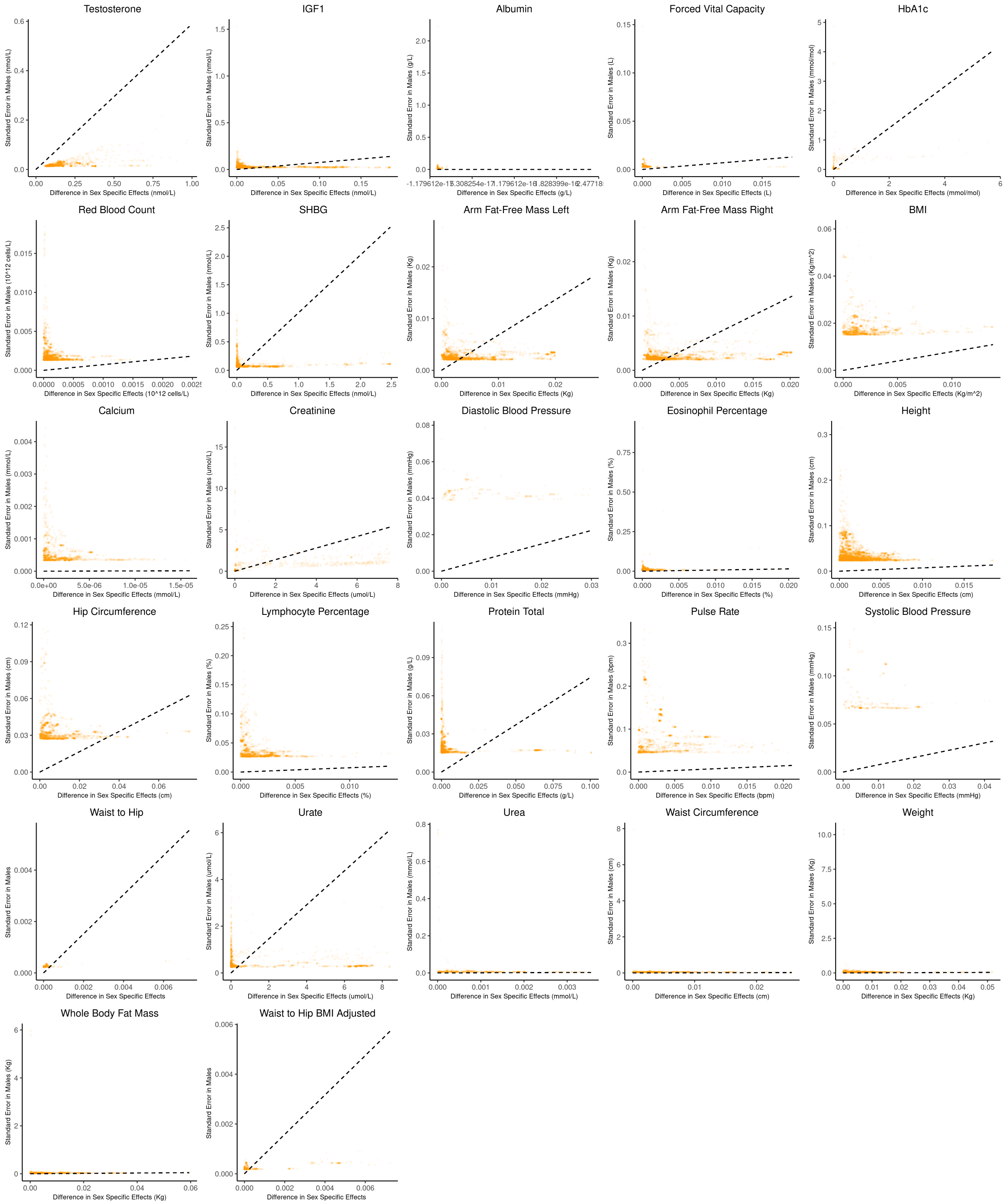

*Supplementary Figure S3: Bias-variance tradeoff at genome-wide significant variants (p-value $< 5 \times 10^{-8}$ in males) in a sex-stratified GWAS. This figure extends Fig. 3B of the main text to all physiological traits analyzed. The x-axis shows the estimated absolute difference between the effect of variants in males and females. The y-axis shows the measured standard error for each variant in males, the focal context here. The dashed line shows the decision boundary for effect estimation in females. The difference in MSE between estimation methods increases linearly with distance from the dashed line, as in **Fig. 2**. If a variant falls above (below) the line, the additive (GxE) estimator has a lower MSE.*

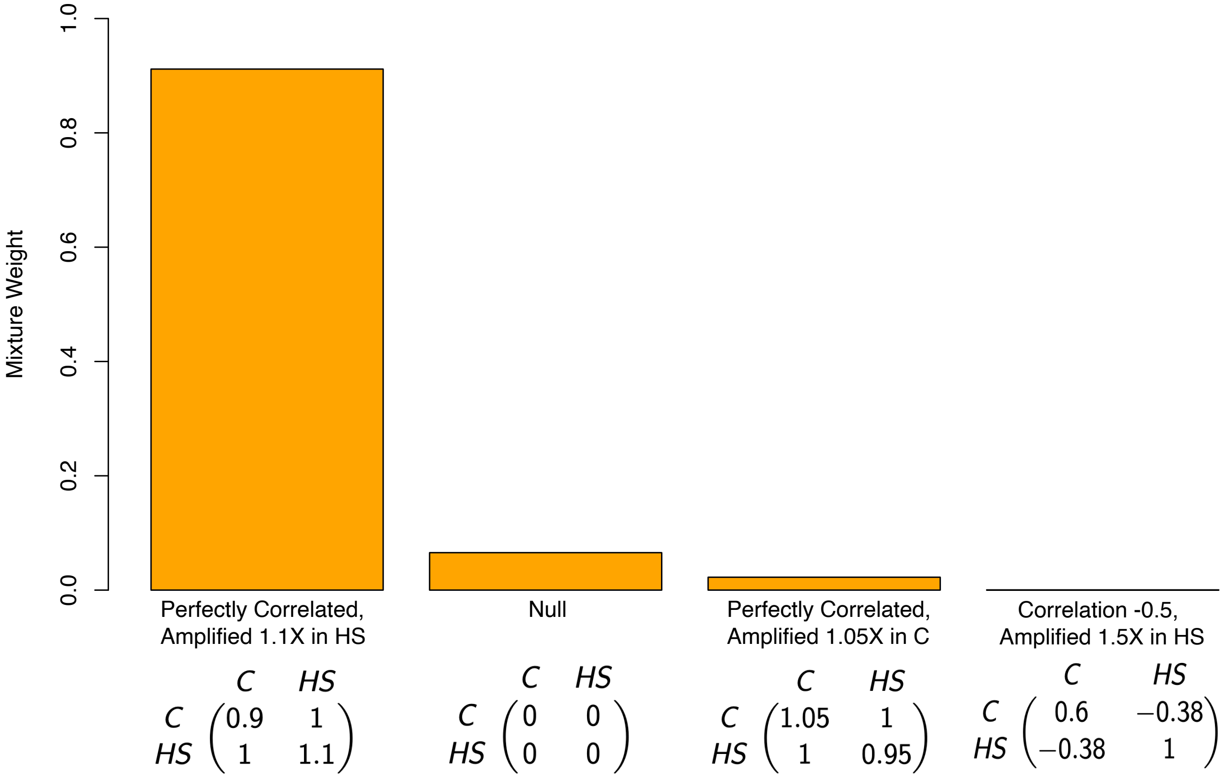

*Supplementary Figure S4: Estimated mixture weights on covariance of true effects in Pallares et al. The x-axis shows (symmetric) variance-covariance matrices for true effects in the high-sugar and control diets. The variance-covariance matrices displayed are the only matrices to which mash assigned non-zero weight (from a much larger set of possible covariance matrices, following a variable selection procedure). Variance-covariance matrices are scaled by a constant for each of interpretability. Abbreviations: C–Control diet; HS–high sugar diet.*

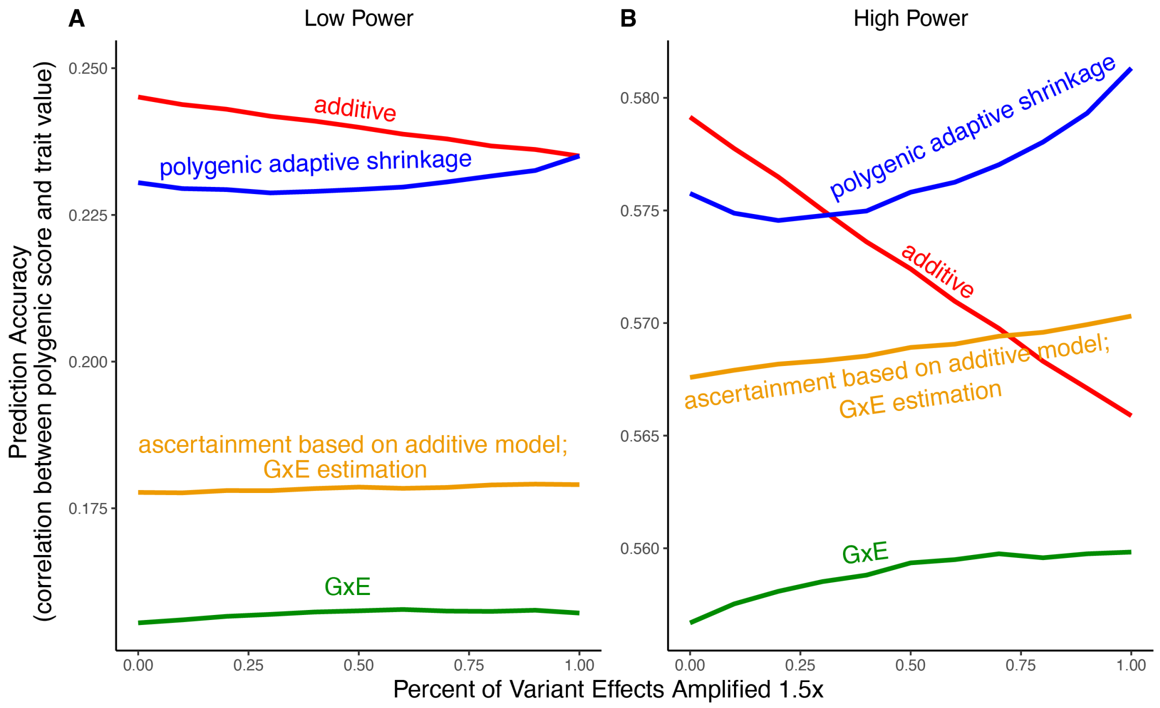

*Supplementary Figure S5: Polygenic score performance for context-aware models with greater amplification than in the main text. This figure uses the same simulation parameters as Fig. 4 of the main text, except that variant effects are amplified* $1.5\times$ *instead of* $1.4\times$*. In each simulation, a GWAS is performed on* $5,000$ *biallelic variants, half of which have no effect in either context. Of the other half, some percent of the variants (indicated on the x-axis) had effects* $1.5\times$ *larger in one of contexts and the remaining SNPs had equal effects in both contexts. The broad sense heritability was set to* $0.4$ *in all simulations. The y-axis shows the average, over* $1,000$ *simulations, of the out-of-sample Pearson correlation between polygenic score and trait value.* ***(A)*** *Results with a GWAS sample size of* $1,000$ *individuals.* ***(B)*** *Results with a GWAS size of* $50,000$ *individuals. We note that unlike in Fig. 5, for amplification percentages greater than or equal to* $75\%$*, the strategy using ascertainment based on the additive model with GxE effect estimation (orange) outperforms the strategy of using the additive model for both tasks (red).*

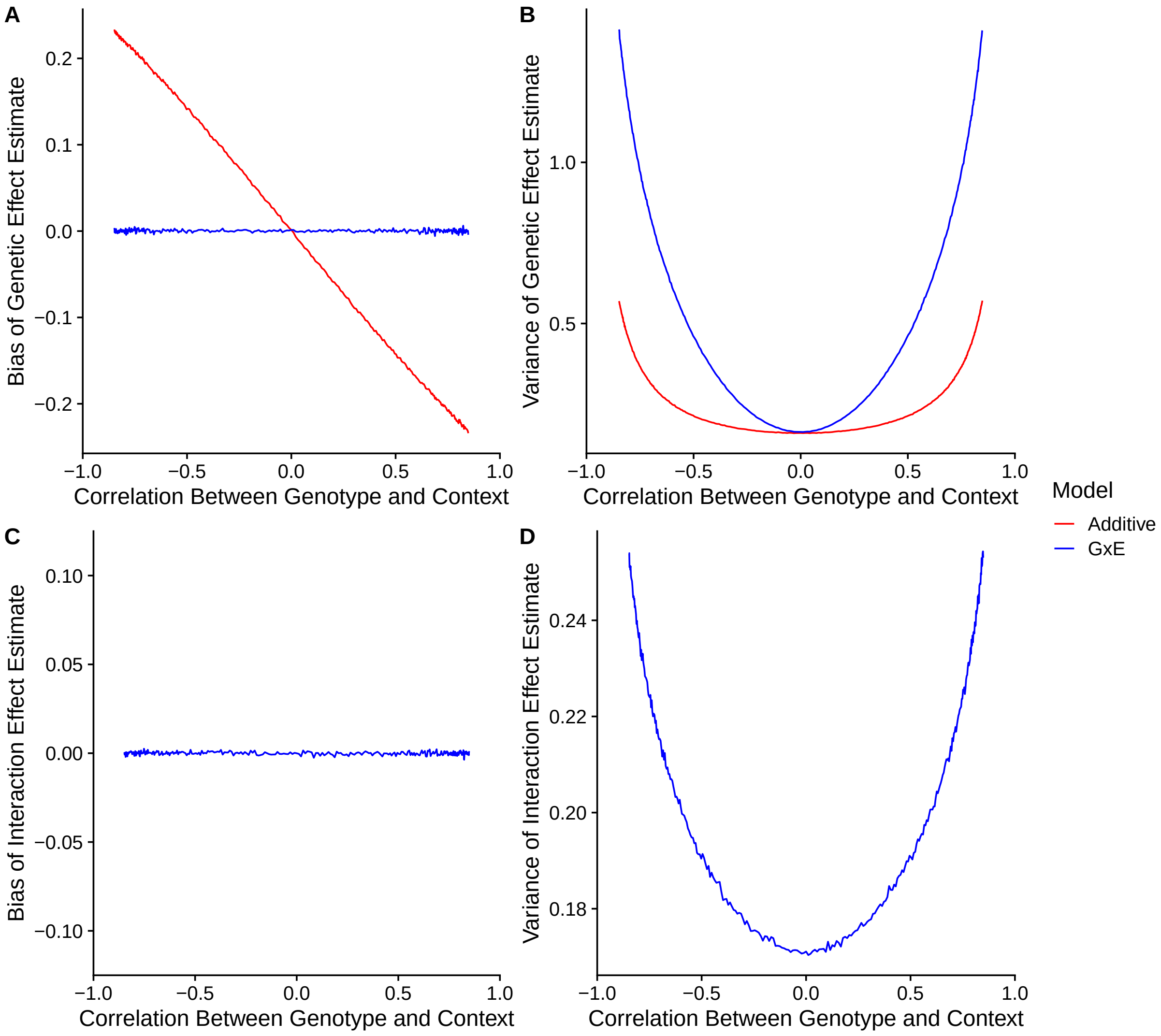

*Supplementary Figure S6: Model performance under correlation between genotype and context.* ***(A)*** *Bias of genetic effect estimate across different values of correlation.* ***(B)*** *Variance of genetic effect estimate across different values of correlation.* ***(C)*** *Bias of term estimating the interaction between genotype and context in an interaction model.* ***(D)*** *Variance of term estimating the interaction between genotype and context in an interaction model.*
